# Construction of a 3-color prism-based TIRF microscope to study the interactions and dynamics of macromolecules

**DOI:** 10.1101/2021.05.10.443430

**Authors:** Max S. Fairlamb, Amy M. Whitaker, Fletcher E. Bain, Maria Spies, Bret D. Freudenthal

## Abstract

Single-molecule total internal reflection fluorescence (TIRF) microscopy allows for real-time visualization of macromolecular dynamics and complex assembly. Prism-based TIRF microscopes (prismTIRF) are relatively simple to operate and can be easily modulated to fit the needs of a wide variety of experimental applications. While building a prismTIRF microscope without expert assistance can pose a significant challenge, the components needed to build a prismTIRF microscope are relatively affordable and, with some guidance, the assembly can be completed by a determined novice. Here, we provide an easy-to-follow guide for the design, assembly, and operation of a 3-color prismTIRF microscope which can be utilized for the study macromolecular complexes, including the multi-component protein-DNA complexes responsible for DNA repair, replication, and transcription. Our hope is that this article can assist laboratories that aspire to implement single-molecule TIRF techniques, and consequently expand the application of this technology to a broader spectrum of scientific questions.

## 1. Introduction

Prism-based single-molecule total internal reflection fluorescence (prismTIRF) microscopy is becoming ever-more popular due to its wide variety of applications. Multiple publications are available which outline the various capabilities of prismTIRF microscopes [1, 2], the rationale behind fluorophore selection [3–5], the general assembly of a 2-color prismTIRF microscope [6–8], and the basics of how to perform such experiments [3, 9–12]. However, even with the currently available guides, building and operating a prismTIRF microscope without expert assistance can be a challenge. With this in mind, we provide here an easy-to-follow detailed guide for designing, assembling, and operating a 3-color prismTIRF microscope which can be utilized for the study of biologically relevant macromolecular assemblies, including protein-DNA complexes.

This guide is broken into four main sections. The first section (**Design**) focuses on the rationale behind selecting certain key components of the prismTIRF microscope, as some components will vary depending on the experimental requirements. The second section (**Construction**) provides a detailed assembly procedure. This procedure is divided into three subsections: the excitation beam path (3.1), which encompasses the components along the excitation path from the lasers to the microscope stage; the microscope stage (3.2), which includes the inverted microscope and the components mounted to the stage; and the emission beam path (3.3), which encompasses the components from the microscope to the detector. The third section (**Operation**) outlines the basics of prismTIRF microscope operation with a primary focus on the acquisition of total internal reflection, alignment of the excitation beams, and the assembly of bead slides and sample chambers. Finally, the fourth section (**Application**) provides pertinent examples of how TIRF microscopes have been utilized to study DNA replication, DNA repair, and DNA transcription.

## 2. Design

### 2.1. Prism-based v.s. Objective-based TIRF

There are two main configurations of TIRF microscopes: objective-based and prism-based. In objective-based configurations, the excitation laser approaches the sample from below the microscope stage and passes through a specialized objective which directs the beam at the appropriate angle to induce TIRF [13, 14]. Because the excitation light approaches from underneath, objective-based configurations allow the top of the sample to be unobstructed and are ideal for imaging open samples such as culture dishes [7, 13]. In prism-based configurations, the excitation laser is typically directed through a quartz prism positioned above the sample (figure 3A). Thus, in prismTIRF the surface of the quartz slide within the sample chamber is imaged, whereas the surface of the glass coverslip is imaged in objective-based configurations. This means that prismTIRF configurations can be used to image samples during active flow, which is challenging to do using an objective-based microscope due to the pressure-induced warping of the coverslip. Additionally, prismTIRF allows for easier manipulation of the excitation beam trajectory and a slightly higher signal-to-noise ratio. Therefore, prismTIRF is well-suited for studying reactions that can be contained within a sample chamber, including interactions between fluorescently labeled macromolecules and surface-tethered substrates. Furthermore, prismTIRF microscopes are relatively simple to assemble and operate, which make them ideal for first-time users. We focus here on the assembly of a prismTIRF microscope (figure 1). Resources regarding the construction of an objective-based TIRF configuration [1, 7, 15], including the cost-effective smfBox [8] which is capable of single-molecule Förster Resonance Energy Transfer (smFRET) experimentation, are available at the associated references.

**Figure 1.**
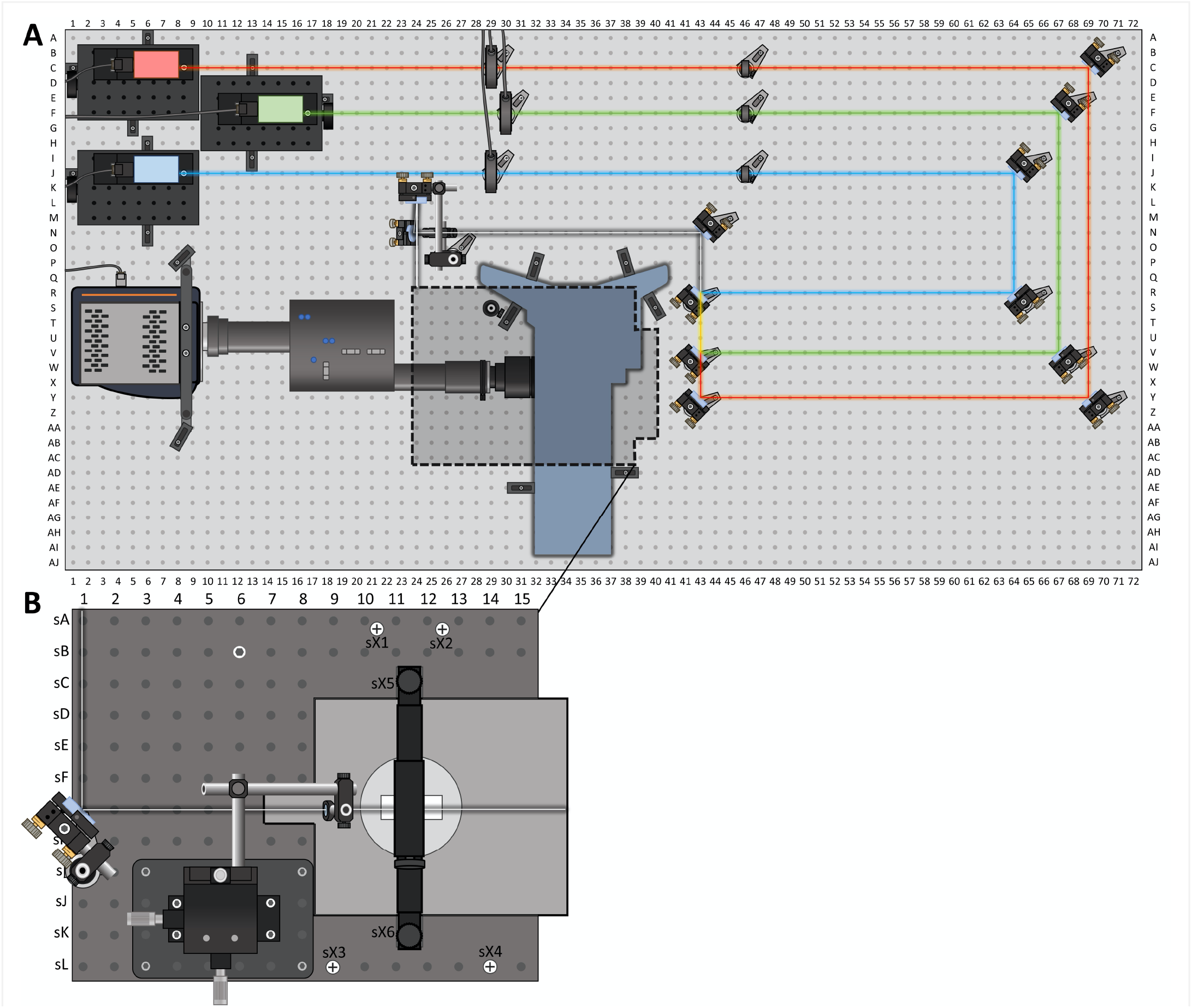
Layout of the prismTIRF microscope. (A) The excitation laser beams (A1-M18) pass through shutters (A28-K31) and clean-up filters (B45-K47) before being leveled with a two-mirror turn (A64-AA72). The beam paths are then merged using two dichroic mirrors (Q41-AA44). The combined beams are redirected (N43) to a periscope that supports two mirrors which elevate the beam to the level of the stage (J22-P28). (B) The microscope stage is fitted with a custom aluminum breadboard (sA1-sL15) that is attached via the front (sX3 and sX4) and rear (sX1 and sX2) stage adapter pieces, and stabilized by a support post that spans between sB6 of the stage breadboard and S29 of the optical table. The beam is reflected from the top mirror of the periscope to the stage mirror (sG1) and passes through a plano-convex lens (sG9) attached to an XYZ linear stage (sI3-sL8). The beam then passes through a quartz prism (sG11-sG12) positioned on top of the sample chamber. The prism is secured in place with an overhead arm mounted to the breadboard (sX5-sX6). The emission path passes through the inverted microscope (P28-AJ40) and enters the Optosplit III (R10-Y29), which is attached to the left port via the C-mount camera adapter (V29-Y32). Dichroic mirrors and emission filters within the Optosplit III (V18-X22) partition the emission from each fluorophore before the image is projected onto the detector (O1-AB10).

### 2.2. Fluorescent Labels and Excitation Lasers

One of the first steps in designing a prismTIRF microscope is to decide on the fluorophores that will be utilized during experimentation. This will dictate the choice of lasers, mirrors and filters within the excitation path, and mirrors and filters within the emission path. For experiments where different colored fluorophores will be simultaneously excited and imaged, it is ideal to use a series of fluorophores which have spectrally distinct excitation and emission spectra. This ensures that each excitation laser only excites the intended fluorophores and that the emission from each type of fluorophore can be easily distinguished (figure 2). The SearchLight Spectra Viewer program provided by Semrock is a good resource for deciphering overlap of excitation and emission spectra (https://searchlight.semrock.com) [16]. The brightness, stability, and Förster resonance energy transfer (FRET) characteristics of fluorophores are also important attributes to consider, and excellent resources are available which provide extensive details in this regard [1, 4, 17].

**Figure 2.**
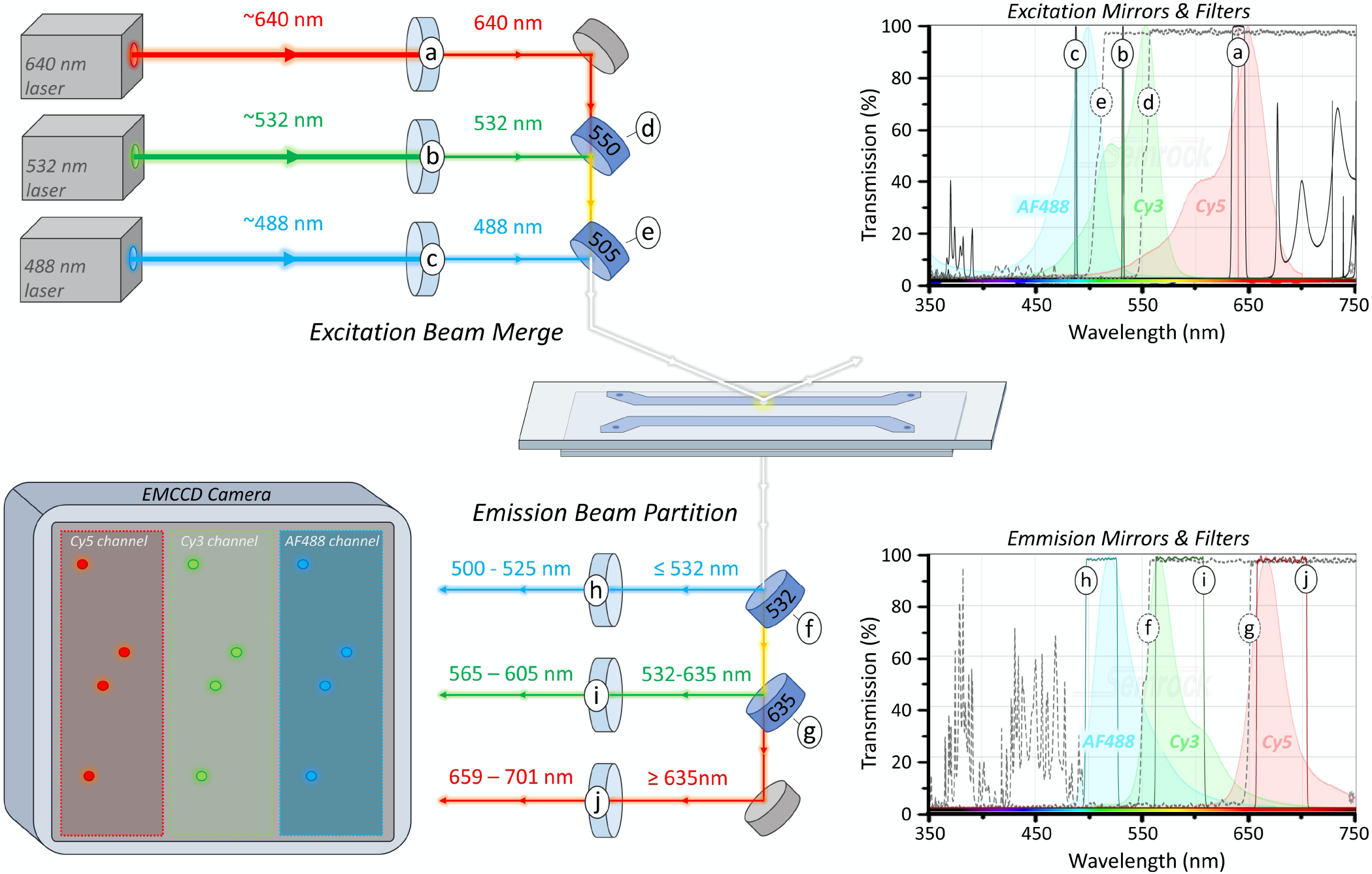
Dichroic mirror and filter scheme. Clean-up filters (a-c) block unwanted excitation energy from reaching the sample. The 640 nm beam passes through the back of the 550 nm long-pass dichroic mirror (d) while the 532 nm beam is reflected from the front, merging the path of the two beams. To combine the 488 nm beam into the laser path, the 640/532 nm beam is passed through the back of the 505 nm long-pass dichroic mirror (e) while the 488 nm beam is reflected from the front. Emission from AF488 is partitioned with a 532 nm long-pass dichroic mirror (f), while the Cy3 and Cy5 emission are seperated with a 635 nm long-pass dichroic mirror (g). Emission filters (h-j) are then used to eliminate bandwidths of light not pertaining to the intended fluorophore. The resulting emission channels are projected onto the EMCCD camera such that they fill the detector space while not overlapping.

Three fluorophores were selected to be utilized in the prismTIRF microscope described here: Cyanine-5 (Cy5), Cyanine-3 (Cy3), and Alexa Fluor 488 (AF488). These are some of the more commonly used fluorophores for single-molecule TIRF experiments due to their stability, brightness, FRET capacity, and well separated excitation/emission spectra (figure 2). However, the infrared fluorophore Cy7 [5], environmentally sensitive MB543 [18], and the many Alexa Fluor variants have also been used with great success. Although this guide is written specifically with these fluorophores in mind, the rationale behind component selection described below applies to any set of chosen fluorophores.

The ideal excitation laser for a particular fluorophore has a wavelength which maximizes excitation of the intended fluorophore while minimizing the excitation of others. The excitation spectra of AF488, Cy3, and Cy5 and the lasers chosen to excite them are shown in figure 2. Although the excitation maxima of Cy3 is 554 nm, the optimal excitation wavelength in this case is 532 nm, since it is minimally absorbed by AF488 and Cy5 (figure 2, b). Similarly, 488 nm is the optimal excitation wavelength for AF488 excitation because it minimizes the excitation of Cy3 (figure 2, c). Although Cy3 absorbs 488 nm light to some degree, the Cy3 emission will be relatively dim compared to AF488 and can be accounted for during data analysis [5]. Typically, lasers rated with a maximum power-level of at least 50-100 mW are sufficient for TIRF imaging [9].

### 2.3. Long-Pass Dichroic Mirrors

To achieve an overlapping excitation field of all three lasers at the region being imaged, the paths of the three beams are merged before reaching the sample (figure 2, d-e). Additionally, since the detector cannot differentiate between wavelengths of light, the emission from each of the three fluorophores must be partitioned and projected onto separate regions of the detector (figure 2, f-g). Both the merging of the excitation beams and the partitioning of emission light are achieved using long-pass dichroic mirrors, which reflect light below a cut-off wavelength while being transmissive to wavelengths of light above the cut-off [19]. In this section, we discuss the rationale behind selecting dichroic cut-offs and the mirror quality requirements at each position.

Merging the three excitation beams is done using two long-pass dichroic mirrors (figure 2, d-e). The first dichroic mirror is used to merge the two longest excitation beams and is positioned such that the longer wavelength beam (640 nm) passes through the back of the mirror, while the shorter wavelength beam (532 nm) reflects from the front. In order for the mirror to be transmissive for the longer wavelength and reflective of the shorter wavelength beam, a cut-off between the two wavelengths is selected (550 nm) (figure 2, d). The second dichroic mirror serves to merge the shortest wavelength beam (488 nm) with the other two and requires a cut-off wavelength between the shortest and second-shortest beam (488/532 nm; 505 nm) (figure 2, e). Again, the two longer wavelength beams should pass through the back of the mirror and align with the path of the reflected beam, merging the three excitation beams.

Emissions from the three fluorophores are partitioned using two dichroic mirrors positioned within an Optosplit III filter cube (figure 2, f-g). The ideal cut-off for these dichroic mirrors will reflect as much emission from the intended fluorophore as possible while being transmissive to the emission of the other fluorophore(s). For example, AF488 emission can range from ~490 nm to ~625 nm and partially overlaps with the emission spectra of Cy3 (figure 2, f). To reflect the most amount of AF488 emission while minimizing the reflection of Cy3 and Cy5 emission, a dichroic mirror with a cut-off of 532 nm is used. The second dichroic mirror within the Optosplit III is used to reflect emission of the fluorophore with the second shortest emission wavelength (Cy3) (figure 2, g). In this case a cut-off of 635 nm is selected, which will reflect nearly all Cy3 emission (532-635 nm) while being transmissive to the majority of Cy5 emission (635-750 nm). After the emissions are separated, bleed over emission from fluorophores with overlapping emission spectra are attenuated using emission bandpass filters (see section 2.4) (figure 2, h-j).

Finally, using mirrors with the appropriate flatness is critical to acquiring clear images. When images are reflected off of an insufficiently flat mirror, the location of the focal plane can shift along the optical axis and the focus spot can be distorted causing the image to become blurry [20]. The emission light that is projected onto the detector is a two-dimensional image and contains pertinent X-Y information. Thus, mirrors within the emission path must be of high enough quality to minimize image distortion. We have found that within the emission path of our set up, 1 mm thick BrightLine® dichroic mirrors with a reflected wavefront error of < 1λ P-V @ 632.8 nm are sufficient to yield a clear image. However, we suggest contacting the supplier directly to discuss their recommendations before purchasing. Conversely, excitation lasers do not contain pertinent X-Y information, so the quality of mirrors within the excitation path is of less consequence, as long as the wavefront error is < 6λ P-V @ 632.8 nm [20].

### 2.4. Bandpass Filters

Bandpass filters are only transmissive for certain bandwidths of light and are used in both the excitation and emission paths to eliminate undesired wavelengths of light. In the excitation path, laser clean-up filters with narrow transmissive bands are used to exclude any erroneous excitation energy from reaching the sample (figure 2, a-c). This helps prevent indiscriminate fluorophore excitation and minimizes background noise. Additionally, after the emission from each fluorophore is partitioned, selective bandpass filters are used to block excitation light and bleed over emission from other fluorophores from reaching the detector (figure 2, h-j). In some cases, it is possible to select emission filters which exclude nearly all emissions from unintended fluorophores, such as the emission filters used here for AF488 and Cy5 (figure 2, h and j). However, for fluorophores with an emission spectra which is overlapped by another fluorophore, such as Cy3, a filter should be selected which has a transmissive bandwidth that maximizes the percentage of intended light over erroneous light (figure 2, i). This can be estimated using the Optimization Calculator within the SearchLight spectral viewer provided by Semrock [16]. Additionally, we suggest contacting the supplier for recommendations to select to optimal arrangement of mirrors and filters. After selecting the optimal dichroic mirrors and band-pass filters to acquire the best possible emission separation from the three fluorophores, the remaining bleed through can be corrected for during analysis [5, 21].

## 3. Construction

### 3.1 Excitation Beam Path

#### 3.1.1. Safety Considerations

The lasers used in this setup are rated Class 3B, which can heat skin and materials but are not considered a burn hazard. However, at 5 to 50 milliwatts of power, a Class 3B laser does pose a moderate risk of eye injury, especially if the laser light remains on one spot on the retina long enough for heat to build up to injurious levels. Thus, it’s important to take measures to prevent direct eye exposure, especially during construction when there is a high risk of inadvertent eye exposure. Laser safety goggles specific for the wavelengths of light being used should be worn during construction while the excitation lasers are active (https://lasersafetyindustries.com). In order to see the laser beam while working, the goggles should have an OD of 5-6, which is high enough to protect the eyes but will not block 100% of the laser’s light. Additionally, black out curtains or a foam board perimeter should be installed to contain reflected laser light within the set-up during everyday operation. Finally, in most cases the Class B lasers must be registered with the institutional laser safety and the appropriate safety warning signage will need to be posted.

#### 3.1.2. Optical Table and Operating Space

The optical table (table 1, #1-2) is the foundation of the prismTIRF microscope (supplemental figure 1A). To provide ample room for the mounting of all components, we recommended using a table that is 3’ by 6’ and fitted with ¼”-20 NC tap holes on a 1” grid. However, the layout depicted in figure 1 could be accommodated to fit on a smaller table by reducing the distance between components in the excitation beam path. The main constraint regarding space is the fixed components within the emission path (i.e. inverted microscope, Optosplit III, and EMCCD). Thus, if space is limited, we suggest using a low-profile alternative to the Optosplit III. The table should be constructed in a low-traffic isolated area to minimize vibrations and potential table bumps which can disrupt image quality and mirror alignment. Additionally, the location must be capable of being completely darkened during microscope operation and, if needed, should be surrounded by black out curtains to reduce aberrant light noise and minimize unintended laser exposure to non-users. Over time, dust particles can settle on the optical components which will require regular cleaning. Thus, it can be beneficial to minimize the amount of dust and particles in the air by running an air filtration device (table 1, #80) nearby and/or installing air filters on nearby vents. Finally, it is recommended to assemble an overhead table shelf (table 1, #3) over the optical table to allow off-table storage of power supplies and provide easy access to the laser and shutter controllers.

#### 3.1.3. Excitation Lasers, Shutters and Clean-Up Filters

The three-color prismTIRF microscope described here is fitted with 488 nm, 532 nm, and 640 nm excitation lasers (table 1, #16-18) allowing for the simultaneous excitation of AF488, Cy3, and Cy5 fluorophores (supplemental figure 1B). To ensure thermal stability of the lasers, each can optionally be mounted to a heat sink (table 1, #19) which will minimize temperature fluctuations during operation. The three laser/heat sink assemblies are attached to vertical translation stages (table 1, #20) and secured in place using L-shaped clamps (table 1, #11). This allows for coarse adjustment to the height of the beam off the table surface. As shown in figure 1, each laser assembly is positioned such that the 640 nm, 532 nm, and 488 nm beams travel respectively over the “C”, “F”, and “J” rows of tapped holes in the table. Using the beam height measurement tool (table 1, #70), the height of each vertical translation stage is adjusted such that the 640 nm, 532 nm, and 488 nm beams are 8” off the table surface when they reach the C69, F67, and J64 holes, respectively (figure 1). Finally, each excitation laser is wired to an adjustable power source (table 1, #21) positioned on the overhead shelf (supplemental figure 1C). Optionally, the adjustable power source can be connected to a nearby computer and controlled using the Coherent Connection software provided by the manufacturer.

The excitation lasers require time to warm up and shut down (~5 minutes), so shutters are incorporated in the beginning of each excitation laser’s path to allow for quick and independent toggling of the excitation beams (supplemental figure 1B). Each shutter (table 1, #22) is attached to a 6” post and an adjustable post holder and secured to the table using a pedestal base adaptor and slotted clamping fork such that the beam passes through the center of the aperture (figure 1 - C29, F30, J29). Optionally, these shutters can be outfitted with ‘Electronic Sync’ that enables computer-controlled shutter operation allowing for Alternating Laser Excitation (ALEX) experiments [22, 23]. Lastly, each shutter is hardwired to a shutter driver switch box (table 1, #23) positioned on the overhead table shelf above the inverted microscope (supplemental figure 1C).

At times, excitation lasers can emit erroneous energy and unintended wavelengths of light. Clean-up filters eliminate any of this undesired excitation energy from reaching the sample (figure 2, a-c). Each wavelength-specific clean-up filter (table 1, #24-26) is secured inside a fixed lens mount (table 1, #14) using a 1/2” spanner wrench (table 1, #63) and attached to a 6” post, adjustable post holder, and pedestal base adapter. The clean-up filter assembly is then mounted to the optical table immediately following the shutters using a slotted clamping fork such that the beam passes through the center of the filter (figure 1 - C46, F46, J46) (supplemental figure 1D).

#### 3.1.4. Leveling and Merging the Laser Beam Paths

After the beams pass through the shutters and clean up filters, they each are reflected from two broadband mirrors that together turn the beams 180°. This two-mirror turn is a tactic that allows for the trajectory of each beam to be fine-tuned, such that they travel level with the surface of the table. To set up the two-mirror turn, six mirror assemblies (two for each beam) containing broad band mirrors (table 1, #27) are constructed using clear-edge mirror mounts, 6” posts, adjustable post holders, and pedestal adaptors (table 1, #7-10, and #13). The mirror assemblies are then mounted such that the beams strike the center of the mirrors directly above the tapped hole of the table as indicated in figure 1 (C69, Y69, F67, V67, J64, and R64) and are reflected at a 90° angle.

Leveling each beam requires the use of two irises positioned after the second mirror (supplemental figure 2). Each iris (table 1, #68) is attached to a 6” post, adjustable post holder, and a ¼”-20 bolt, and adjusted such that the center of aperture is 8” off the surface of the table using a ruler or a beam height measuring tool (supplemental figure 2A). A slip-on post collar (table 1, #69) can then be attached to the iris assembly which will allow the post to be rotated within the adjustable post adapter without altering the height of the iris. Starting with the shortest wavelength beam path, the two 8” irises are then mounted directly into holes of the table, via the ¼-20 bolt, just beyond the second mirror, (R67 and R46 for 488 nm beam) (supplemental figure 2B-C). Optionally, if the microscope has not already been placed on the table, the second iris can be placed at the edge of the table (R1, V1, and Y1) to maximize the distance between the two irises, which will increase the precision of the laser leveling. After leveling the shortest wavelength beam (see below) the irises are moved to the next longest wavelength beam to level that beam, and again for the longest wavelength beam (V65 and V46 for the 532 nm beam, and Y62 and Y46 for the 640 nm beam).

With the two irises in place, the beam can be ‘walked’ into position by adjusting the first and second mirrors iteratively. To start, the first mirror is adjusted so that the beam reflects off the second mirror and passes through the very center of the nearest iris (supplemental figure 2B). Then, the aperture of the nearest iris is opened slightly, and the second mirror is adjusted so the beam passes through the center of the far iris (supplemental figure 2C). This process is repeated, alternating between adjusting the first mirror so the beam strikes the center of the near iris, and adjusting the second mirror so the beam strikes the far iris, until the beam passes cleanly through the very center of both irises. With the leveling complete for the shortest wavelength beam, the irises are moved into the path of the next longest wavelength beam, and the leveling process is repeated. After all three beams are leveled, each beam will be on a level plane 8” off the table surface. For additional help, video tutorials describing the process of walking a beam into alignment can be found on the Thorlabs website (https://www.thorlabs.com/).

The beam paths are combined using a broadband mirror and two long-pass dichroic mirrors (figure 2, d-e) (figure 5B). To set up the beam merging mirrors, the two 8” tall irises are first mounted directly into the P43 and A43 holes (figure 1). A mirror assembly (table 1, #7-10, and #13) containing the 505 nm cut-off dichroic mirror (table 1, #28) is then placed on the table such that the 488 nm beam reflects at a 90° angle off the front face of the mirror over the R43 hole and passes through the center of the two irises. After the 505 nm mirror is in position, a mirror assembly (table 1, #7-10, and #13) containing the 550 nm cut-off dichroic mirror (table 1, #29) is then placed on the table such that the 532 nm beam reflects at a 90° angle off the front face of the mirror over the V43 hole, passes through the back of the 505 nm dichroic mirror, and passes through the center of the two irises. Finally, a mirror assembly (table 1, #7-10, and #13) containing a broadband mirror (table 1, #27) is placed on the table such that the 640 nm beam strikes the center of the mirror over the Y43 hole, is reflected at a 90° angle, and passes through the back of the 550 nm mirror, 505 nm mirror, and both irises. When all three beams pass through the center of both irises, the beams have been successfully merged. Note that following the order described above for incorporation of the mirrors will make this step significantly easier, as chromatic aberrations in the dichroic mirrors could alter the trajectory of the transmitted beam(s).

#### 3.1.5. Beam to Stage Path

After the excitation beams are merged, they are directed to the stage of the inverted microscope using a series of broadband mirrors (supplemental figure 3A). Note that from this point forward, only one of the three beams should be on while setting up the remainder of the excitation beam path. We recommend using the beam which will be used most often during operation to establish this path and/or the mid-wavelength beam. This is because the other beams will require periodical realignment to match the beam used to establish the excitation path (section 4.2), but the path of the established beam will remain unchanged after construction. Here, the mid-wavelength beam (532 nm) was used to set up the remainder of the excitation path.

A broadband mirror at N43 is used to direct the beam behind the microscope and it is positioned such that the beam is reflected through the center of two 8” tall irises mounted into the N41 and N14 holes (figure 1). The beam is then elevated to the height of the microscope stage using two broadband mirrors attached to an 18” periscope (supplemental figure 3B-C). The periscope is assembled from three 6” optical posts and secured to the table near the O26 hole using a pedestal base adaptor and a slotted clamping fork. The two broadband mirrors are mounted within clear-edge mounts, attached to 2” posts, and then each is secured to the 18” post using two right-angle clamps and a 4” post. The bottom mirror is positioned such that the beam is directed 90° vertically, parallel with the 18” post, and the top mirror is positioned to redirect the beam 90° horizontally toward the front edge of the table. The exact positioning of the top mirror will be determined in section 3.2.2, after the inverted microscope is assembled. An alternative option to the periscope is to use 30 mm cage optics. Although the cage optics will incur an additional cost, it will save time in aligning the excitation beam to the stage.

### 3.2. Microscope Stage

#### 3.2.1. Inverted Microscope and Stage Breadboard

The inverted microscope is placed on the table with the front bottom edge of the microscope sitting roughly 2” from edge of the table and centered between the 38^th^ and 39^th^ column of tapped holes (figure 1, supplemental figure 1). Securing the microscope in place are six L-Shaped clamps at holes AD42, AD35, S43, S34, P41, and P37. Once the inverted microscope is secured in place, the microscope components (Table 1, #30-39) can be assembled according to the IX73 user manual.

A series of components mounted to the microscope stage are used to direct the excitation beam into the sample chamber. To secure these components to the stage, a custom-made stage breadboard (table 1, #42) with ¼”-20 NC tapped holes in a 1” grid is attached to the top of the inverted microscope. Before the stage breadboard can be attached, two custom-made stage adapters (table 1, #40-41) are secured to top of the inverted microscope (supplemental figure 4). The stage breadboard can then be bolted to the microscope via the stage adapters at position sX1-sX4 (table 1, #66-67) (figure 1, supplemental figure 4A and C). These custom components can be commissioned from a local machine shop, using the design files provided in the supplemental files. Finally, to provide additional stability to the stage breadboard, a support beam built from two 6” posts and an adjustable post holder is mounted between the optical table (S34) and the breadboard stage (sB6) (supplemental figure 4C).

#### 3.2.2. TIR Angle and the Stage Mirror

In prismTIRF microscopy, the excitation beam is directed into the sample chamber at a particular incident angle which results in the beam being totally reflected from the slide/aqueous sample interface (figure 3A) [13, 24, 25]. This total internal reflection (TIR) generates an evanescent wave that penetrates ~100-200 nm into the sample chamber, only exciting the fluorophores located within this thin band [14]. The penetration depth of the evanescent wave depends on the incident angle of the beam, the refractive index (RI) of the two mediums at the interface (i.e. the quartz slide and aqueous sample), and the wave-length of the beam [26, 27]. Evanescent waves produced from longer wavelength beams and smaller incident angles will penetrate deeper into the sample. For a quick tutorial on TIR angles and evanescent wave penetration depths, we recommend referencing the evanescent wave penetration depth java tutorial provided by Olympus [28]. In most cases, simply knowing that the system is in the TIRF imaging mode and having a rough estimate of the penetration depth is adequate. However, if quantitative measurements of the penetration depth are desired, there are ways to determine these values [26, 27].

**Figure 3.**
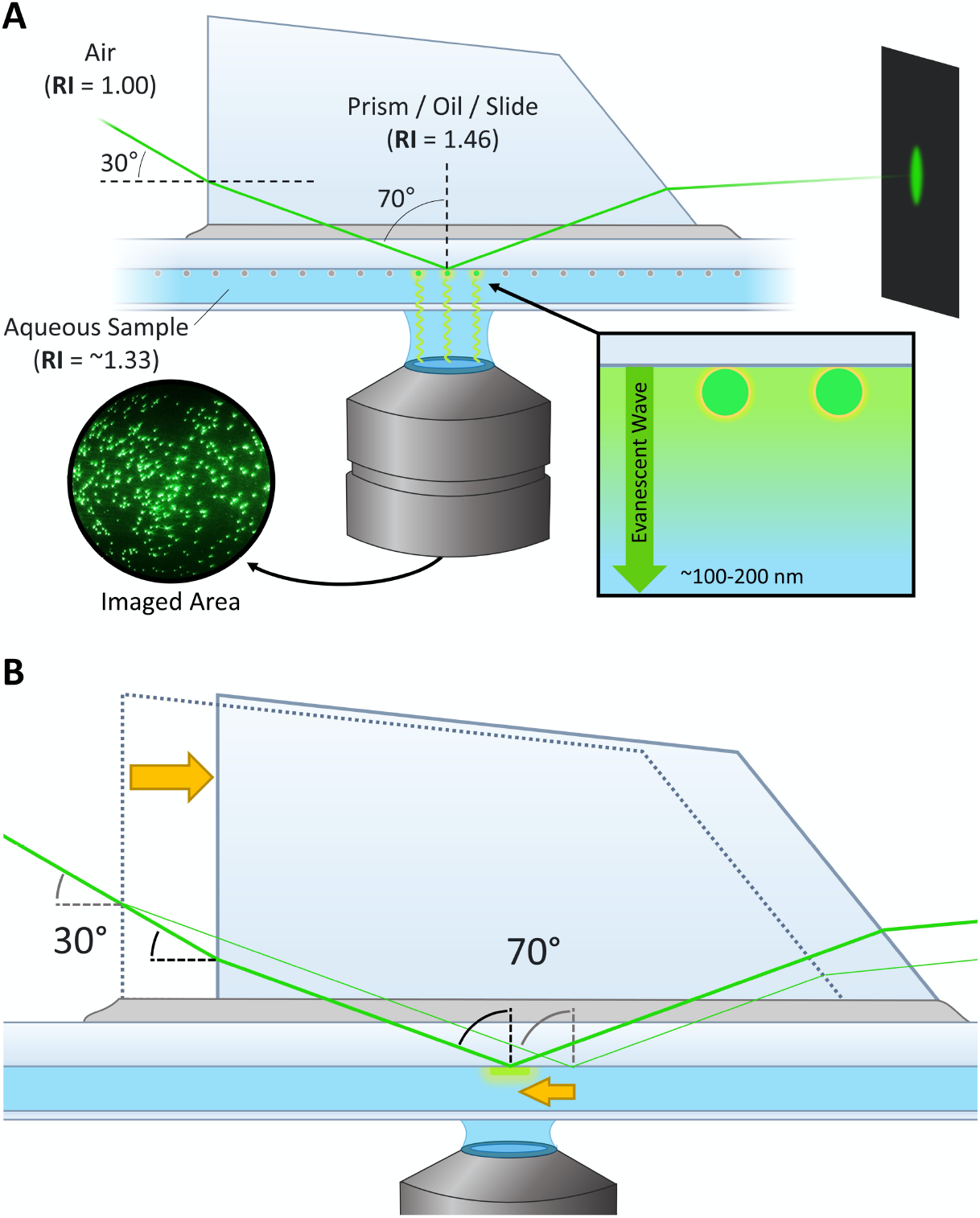
Trajectory of excitation beam through the prism. (A) The excitation beam enters the front face of the quartz prism at an incident angle of ~30°. The difference in refractive index between the air and quartz results in a 20° angle of refraction. The quartz prism, oil, and quartz slide have similar refractive indeces which minimizes further refraction of the beam. The beam approaches the interface of the quartz slide and aqueous sample at a ~70° incident angle, resulting in total internal reflection of the beam and generation of an evanescent wave which emanates 100-200 nm into the sample. When TIRF is achieved at the surface of the quartz slide, the reflected beam should produce a symmetric circle on the far wall. (B) The prism is used to adjust the trajectory of the excitation beam. Sliding the prism horizontally alters the height at which the beam enters the prism, which shifts where the beam strikes within the sample chamber while maintaining the proper 70° incident angle.

To induce TIR at the interface of the quartz slide (RI = 1.46) and aqueous sample within the chamber (RI = ~ 1.33), the incident angle of the co-aligned 488 nm, 532 nm, and 640 nm beams should be between 68-71° (figure 3A) [6, 14, 26, 27]. At 70°, the 488 nm, 532 nm, and 640 nm beams will generate evanescent waves which penetrate roughly 105 nm, 114 nm, and 138 nm into the sample, respectively [14]. Due to the difference in RI between the air (RI = 1.0) and quartz prism (RI = 1.46), the beam will refract upon penetrating the quartz prism before reaching the slide/sample interface. Thus, the beam should enter the quartz prism at a 28-33° incident angle to induce the 68-71° incident angle at the slide/sample interface.

A broadband mirror positioned on the stage breadboard is used to direct the coaligned excitation beam into the sample chamber (supplemental figure 5). The height at which the beam should strike the stage mirror to induce the 30° incident angle at the prism face is dependent on the horizontal distance between where the beam strikes the mirror and the prism face. In this particular set-up, the beam should traverse directly over the sA1-sF1 holes of the stage breadboard and reflect off the stage mirror directly above sG1, which is approximately 26.5 cm (10-7/16”) from where the prism will be positioned (supplemental figure 5A). Considering the surface of the stage breadboard is ~2 cm (25/32”) above the microscope stage, the beam should strike the stage mirror 13.5 cm (5-5/16”) above the surface of the stage breadboard, totaling 15.5 cm (5-29/32”) of vertical difference to induce a 30° incident angle at the prism face.

To set up the stage mirror, two irises with centers at 13.5 cm (5-5/16”) high are placed in the sA1 and sL1 holes of the stage breadboard, and the top mirror on the periscope is adjusted such that the beam travels through the center of both irises (supplemental figure 5B). This will require multiple iterative adjustments to both the bottom and top mirror of the periscope to achieve the proper beam trajectory. When finished, these irises are removed and a broadband mirror is inserted into a clear-edge mirror mount, attached to a 3” post, and combined to a 6” post and 2” post holder using a right-angle clamp (supplemental figure 5C). The stage mirror is then positioned over the sG1 hole in the stage bread-board such that the beam is reflected squarely over the sG2-sG6 holes and strikes the microscope stage between the overhead arm mounting holes (figure 1, supplemental figure 5D). To observe the position where the beam strikes the stage, a glass microscope slide covered with opaque tape can mounted to the stage. To ensure that the beam is traveling squarely over the sG2-sG6, two 6” posts can be inserted into the sG2 and sG6 holes of the stage breadboard. If the beam is square over the holes, it should strike the center of the sG2 post, and after the sG2 post is removed, the beam should also strike the center of the sG6 post. Alternatively, irises can be used instead of 6” posts for a more precise calibration of the beam trajectory.

#### 3.2.3. Focusing Lens and Prism Assembly

Optimal TIRF imaging typically requires making small adjustments to the position and diameter of the illuminated field during operation. These adjustments are made using a plano-convex lens (table 1, #50) and a quartz prism (table 1, #71), which can be used to focus the beam and fine tune the beam trajectory. Before setting up the prism and lens, the objective will need to be adjusted to the proper height. This can be done by placing a bead slide (see section 3.1 below) on the stage and focusing on the inside edge of the adhesive within the chamber (supplemental figure 6A) [7]. After focus is acquired, the slide can be removed from the stage and the prism and lens components can be assembled.

The lens is mounted to an XYZ linear stage which allows for fine adjustments to the focus and trajectory of the excitation beam (supplemental figure 6B). To assemble the XYZ linear stage and lens, a 4 × 6” breadboard (table 1, #51) is first mounted over the sI3-sL8 holes (figure 1) to elevate it slightly off of the stage breadboard. The three micrometer actuators (table 1, #53) are attached to the XYZ linear stage (table 1, #52) and set to the center of their travel range (in this case ~6 mm). The lens (table 1, #50) is then inserted into a fixed lens mount (table 1, #14) such that the flat side will face the objective. A 3” post is attached to the fixed lens mount, a 4” post is attached to the top hole of the XYZ linear stage closest to the objective, and the two are connected using a 6” post and two right-angle clamps (supplemental figure 6B). The posts are then adjusted within the right-angle clamps such that the lens is approximately 5 cm from the objective and the beam is transmitted through the center of the lens and strikes the near edge of the objective’s front lens (supplemental figure 6B - inset). When positioned properly, the beam should produce symmetrical points of light on opposite sides of the objective’s front lens.

The quartz prism is attached to a custom adaptor which allows the prism to be easily manipulated during operation of the microscope (supplemental figure 7A). Similar to the stage breadboard and stage adapter pieces, the components used to support the prism (table 1, #43-45) can be commissioned form a local machine shop. To attach the prism to the prism adaptor, the prism connector piece (table 1, #43) is first bolted to the prism adapter (table 1, #44) using the three prism adapter bolts (table 1, #49). A generic glass slide is then placed on the stage and the prism adapter is secured to the overhead arm (table 1, #45) and mounted to the stage breadboard (supplemental figure 7B). A Kim wipe is laid over the slide, and then the loose prism is placed on the Kim wipe directly underneath the prism adapter. A thin coat of 5-minute epoxy is added to the top of the prism and then the prism adapter is released from the overhead arm and lowered until the connector piece makes contact with the top of the prism (supplemental figure 7C). The prism can be repositioned as needed by gently sliding the Kim wipe. The prism adapter is then locked into place and the glue is allowed to set overnight. Attaching the prism in this way helps to ensure the prism is flat with respect to the slide while secured to the overhead arm.

### 3.3. Emission Beam Path

#### 3.3.1. Optosplit III Emission Splitting and the EMCCD

The combined emissions from each fluorophore must be separated into three channels using dichroic mirrors before being projected onto their respective regions of the detector (figure 2, f-g). This is accomplished using an Optosplit III (table 1, #60), which is connected to the left-side port of the inverted microscope (supplemental figure 8). To attach the Optosplit III to the microscope, the C-Mount camera adapter (table 1, #32) is first attached onto the entry port of the Optosplit III. Then, the left side port clamping screw of the microscope is loosened, the cap is removed, the camera adapter is inserted into the C-Port adapter is secured into the left port of the inverted microscope, using the support foot to support the weight of the Optosplit III body. With the Optosplit III attached, the dichroic mirrors and emission filters used to partition the emission are mounted inside the filter cube and inserted into the Optosplit III housing as described in the user manual.

The final component of the prismTIRF microscope is the detector. In this set-up, an electron multiplying CCD (EMCCD) camera (table 1, #59) is used as the detector device, although other comparable options are available [29, 30]. The EMCCD is attached to a custom aluminum camera mount (table 1, #46) and secured to the table using L-shaped clamps in order align the EMCCD with the exit port of the Optosplit III housing (supplemental figure 8A). After attaching the EMCCD to the Optosplit III, the EMCCD is connected to the computer via USB and can be operated using imaging software such as Micro-Manager (https://micro-manager.org), or the smFRET data acquisition and analysis package provided by TJ Ha and colleagues (https://cplc.illinois.edu/research/tools) [21]. We use the software package provided by the Ha group, although the choice of software is primarily dependent on user preference and the file format requirements of the analysis software being used. Details regarding the configuration of the camera and operation of the software are available in the reference manual provided with the software package. After the EMCCD is attached, the adjustment knobs on the Optosplit III body can be used to adjust the position of each emission channel on the detector. Detailed information regarding the set-up and operation of both the EMCCD and Optosplit III can be found in the manuals provided by the manufacturers.

## 4. Operation

### 4.1. Bead Slides and TIR Acquisition

A bead slide is a mock sample chamber which contains immobilized multi-fluorescent beads (supplemental figure 9) [7, 14]. The bead slide is used to record a short mapping video which is used to calibrate the software that aligns the X-Y coordinates of the partitioned emission channel projections [3]. Additionally, the bead slide is useful for trouble-shooting TIR acquisition. At times, it can be a challenge to find TIR on an experimental slide, especially if the position of the lens and prism are far from the proper position to induce optimal TIRF. However, the fluorescent beads within the bead slide are easy to see and will be visible even if the beam trajectory is near to, but not perfectly at, the proper angle to induce optimal TIRF. Therefore, it can sometimes be helpful to dial in TIR on a bead slide in order to get close to TIR before attempting to image experimental sample chambers.

Before setting up the microscope for TIRF imaging, the prism and slide are thoroughly cleaned using ethanol. This is critical, as even the smallest amount of debris, dried oil, and/or streaks can cause significant issues to the image and TIR quality. A drop of water is placed on the objective and the stage clips are used to secure the bead slide to the stage. The objective is then adjusted so that the seam of the tape or glue within the chamber is in focus (supplemental figure 6A). This will position the objective such that the TIR signal will be in focus once the prism and lens are properly positioned for TIRF imaging. It’s important not to move the objective too far from this position while acquiring TIR, as it is difficult to focus the objective with the prism in place, and the TIR signal will not be visible if the slide/sample interface is out of focus. After the objective is positioned, the 532 nm beam is turned on and, if necessary, the lens micrometer is adjusted such that the beam strikes the left-most edge of the objective’s front lens (supplemental figure 6C). Finally, the stage is adjusted so that the center of the slide is over the objective and a drop of low-fluorescence immersion oil is placed on the slide. The prism is then mounted to the microscope stage and carefully lowered onto the drop of oil (supplemental figure 10).

Acquiring optimal TIR for TIRF imaging involves tuning the trajectory and focus of the excitation beam by making fine adjustments to the positioning of the prism and lens (figure 4). Sliding the prism horizontally over the surface of the slide will alter where the beam strikes within the sample (figure 3B). When both the lens and prism are positioned properly, a dark background with a field of points of light will emerge (figure 4B-D) and the reflected beam should produce a symmetric circle on the far wall (figure 3A). After securing the prism in place, the position of the lens can be adjusted using micrometers in order to center the beam within the imaged area (figure 4E-G) and expand or narrow the illuminated field (figure 4H). It’s recommended to turn off all lasers except the 532 nm laser when acquiring TIR since the other beams will be aligned with respect to the 532 nm beam later on (section 4.2).

**Figure 4.**
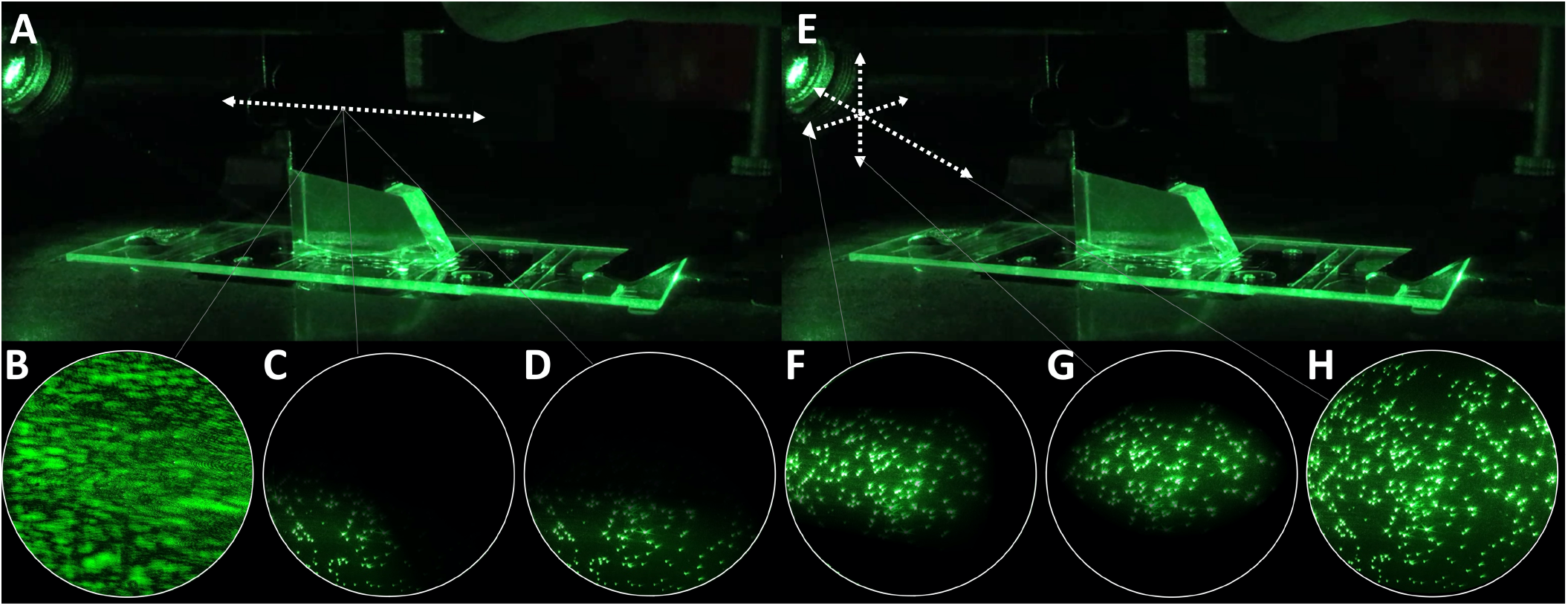
Aquiring TIR. (A) The prism adapter is adjusted horizontally within the overhead arm. To aquire TIR at the imaged area, the 532 nm laser is turned on and the prism adapter is freed from the overhead arm. The prism is then adjusted until the beam illuminates the region of the slide directly above the objective. If the trajectory of the beam is correct, this should cause the hazy light (B) to be replaced with a points of light on a dark background (C). (D) The prism is adjusted until the illuminated field is centered horizontally within the eyepiece and resecured to the overhead arm. (E) The lens micrometers are used to make fine adjustments to the beam trajectory and focus. (F-G) The micrometers are adjusted to center the illuminated field within the imaged area. (H) The illuminated field can be narrowed or expanded to fill the imaged area by moving the lens towards or away from the prism along the beam path, respectively. This is done by incrementally adjusting the lens down and toward the prism, or up and away from the prism, while simultaneously looking through the eyepiece to ensure the illuminated field remains in view.

### 4.2. Excitation Beam Alignment

After the region illuminated by the 532 nm beam is adjusted to fill the center of the imaged area, the other beams (488 nm and 640 nm) may need to be aligned so that they illuminate the same field as the 532 nm beam (figure 5A). If the 488 nm and/or 640 nm beams are out of alignment, the illuminated field will appear off-center within the imaged area. To correct this, the trajectory of the 488 nm and 640 nm beams can be independently adjusted by turning the adjustment knobs on the 505 nm dichroic mirror and broadband mirror used to merge the excitation beams, respectively (figure 5B). When the beams are aligned, the color of the illuminated beads should be homogenous over the imaged area when all lasers are on (figure 5C). Importantly, the trajectory of the 532 nm beam should never be adjusted by turning the adjustment knobs on the 550 nm dichroic mirror, as doing so will cause the path of the merged excitation beam that was established during construction to be lost. The 532 nm beam should only be adjusted using the prism and/or lens micrometers.

**Figure 5.**
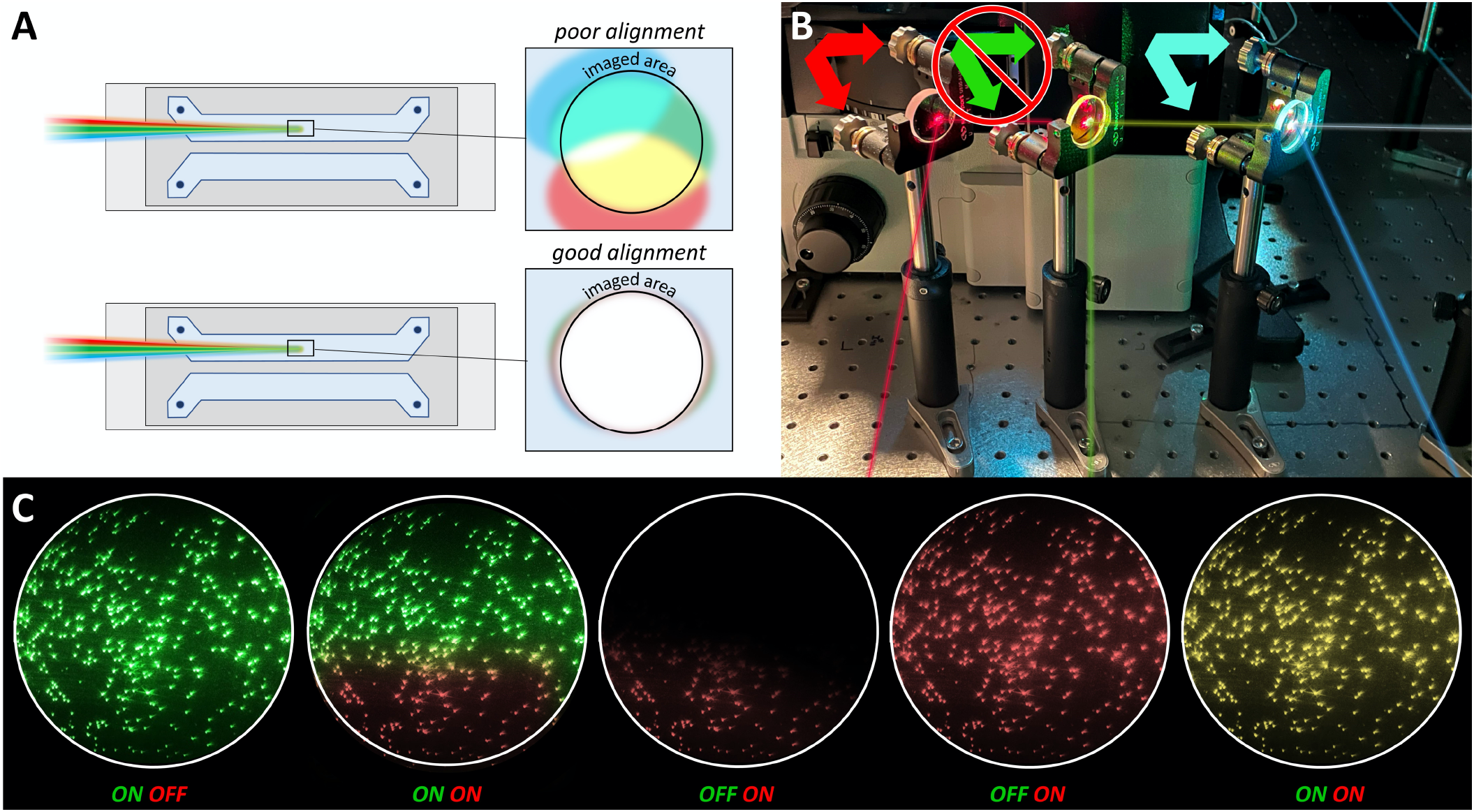
Aligning the excitation beams. (A) The 640 nm and 488 nm beams require alignment if their illuminated fields do not overlap with the centered 532 nm beam. (B) The 640 nm and 488 nm beams can be aligned independently by tuning the adjustment knobs on the broadband mirror at Y43 and the 505 nm cut-off dichroic mirror, respectively. Importantly, the 550 nm cut-off dichroic mirror should not be adjusted, as this will forfeit the original beam path established during construction. (C) The adjustment knobs of the mirror mounts are tuned until the illuminated fields are centered within the imaged area. When the 640 nm and 488 nm beams are properly aligned, the color of the illuminated field will be homogenous when all lasers are on.

### 4.3. Emission Channels and Detector Settings

The emissions are visualized and recorded using the EMCCD, a computer, and an imaging software such as the Single.exe program contained within the smFRET data acquisition and analysis package provided by the TJ Ha group [21]. Here, we provide a brief overview of the operation of the camera and the Single.exe imaging software. More detailed information can be found on the EMCCD manufacturer’s website and in the Single.exe reference manual [21]. To visualize the emission channels, the power button on the back of the EMCCD is turned on and the Single.exe program is opened. The camera is cooled to −80 °C and the PMA recorder window is opened. After confirming that the illuminated field is centered within the eyepiece and the excitation beams are aligned (figure 5C), the light-path is diverted completely to the camera using the light path selector lever on the right side of the inverted microscope. Finally, before activating the camera, the lights in the room are turned off. When imaging a bead slide with Single.exe, the gain should be reduced to ~10 and the ‘Background’ and ‘Data Scaler’ settings should be set to ‘Autoscale’. The 532 nm beam is turned on and, if needed, the laser power should be adjusted such that the bead spots depicted in the imaging software are blue with a slight red center. Bright white spots will saturate the detector result in an unusable mapping file. Note that fine adjustments to the focus of the inverted microscope may be required to sharpen the image. When imaging a sample chamber, the gain should be increased to 299 and the ‘Background’ and ‘Data Scaler’ settings should be adjusted to acquire a clear and well contrasted image. To maintain consistency between data sets, it’s recommended to keep these settings constant between experiments within a dataset.

The position of each emission channel on the detector can be adjusted using the adjustment knobs on the top and side of the Optosplit III (supplemental figure 8A). Detailed information regarding this procedure is outlined in the Optosplit III manual. If three emission channels are to be recorded simultaneously, the size of the channels can be adjusted to each fill a third of the total screen area using the aperture control levers located on the entry port of the Optosplit III (supplemental figure 8A). If only two emission channels are needed for an experiment, a blocker can be easily placed into the Optosplit III housing that will block the undesired emission path and the remaining two channels can be adjusted to each fill half of the detector. The position of each channel should be adjusted similar to the depiction shown in figure 2, with space around the edges of each channel so they do not extend beyond the edges of the detector or overlap with one another.

At the start of each week of experiments, or after making any adjustments to the position of the emission channels, a new mapping file should be recorded. A mapping file is a short 100 frame video of the stationary beads within a bead slide that is used to map the X-Y coordinates of each emission channel [3]. During processing of an experimental data video, the mapping file will be used to define the region of each emission channel so they can be overlayed. This process is explained in detail in the documentation provided with the smFRET data acquisition and analysis package.

### 4.4. Sample Chamber Assembly and Data Collection

The experimental sample chamber contains the fluorescent sample(s) under observation. It consists of a quartz slide with two or more holes drilled into its surface, a glass coverslip, and an adhesive to attach the two together and create a channel between the holes (figure 6) [31]. The quartz slide and coverslip require extensive cleaning and passivation to minimize background and noise, and detailed information regarding this process is readily available [32–34]. After the surfaces are prepared, sample chambers have traditionally been constructed using double stick tape to create flow channels between the holes in the slide and epoxy to seal the ends [32]. Although double-stick tape is effective and is still used successfully in many labs, we have found that using adhesive sheets (table 1, #81) cut with either a razor blade or a cutting machine (e.g. Cricut) to build the flow channels prevents the need of 5-minute epoxy to seal the ends of the channel and significantly reduces the time and technicality of building sample chambers (figure 6 A-H). The cutting machine designs we use can be found in the supplemental files.

**Figure 6.**
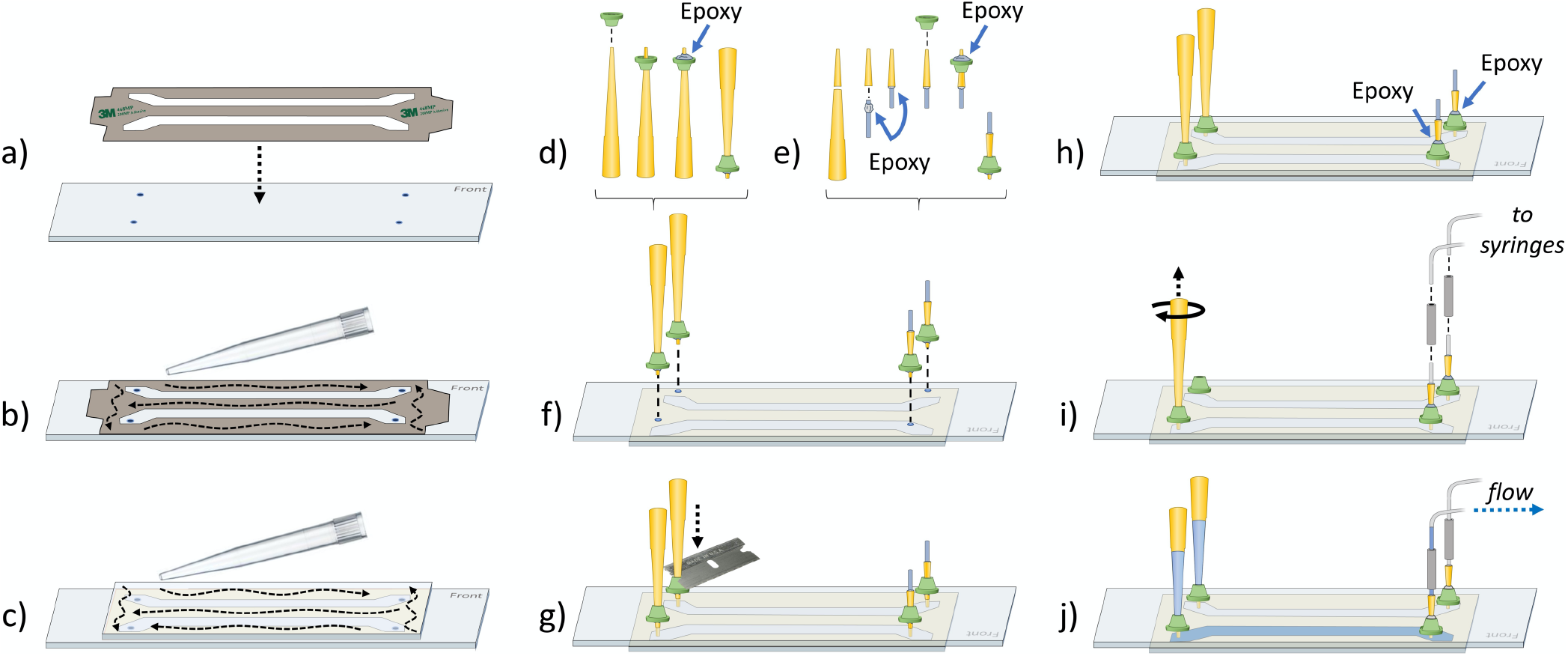
Building sample chambers. (a) A sheet of double-sided adhesive is cut to create flow chambers between the drilled holes of the quartz slide. The quartz slide is placed on a clean surface PEG-side up. One side of the adhesive backing is removed and the adhesive is positioned onto the slide. (b) A pipette tip is pressed against the back of the adhesive to adhere it to the slide. (c) The second backing of the adhesive is removed and a coverslip is placed over the adhesive, PEG-side down. A razor is used to remove any excess adhesive surrounding the coverslip and a pipette tip is used to press on the coverslip to secure the bond. The sample chamber is then flipped over so the holes are exposed. (d-g) The entry and exit ports are installed one at a time. (d) Entry ports are installed by sliding a 3D printed port onto the end of a 200 μL tip. A smear of 5-minute epoxy is applied to the bottom of the port carefully to avoid getting glue on the very end of the tip. (f) The tip is inverted and inserted into a hole on the sample chamber. (g) While using a finger to put pressure on the top of the tip to keep it in the hole, the side-notch of a razor blade is used to push the port down firmly to the slide surface. This is repeated to install the second entry port. (e) To install an exit port, the top ¾” is severed from a 200 μL tip (table 1, #79) using a razor blade. A ring of glue is applied to one end of a 1” long piece of ETFE tubing (table 1, #78) and inserted firmly into the shortened tip. Batches of these tip/tubing assemblies are made in advance to save time. Exit ports are then installed following the same procedure as the entry ports (e-g). (h) Epoxy is applied to the top of each exit port to ensure an air-tight seal. The 5-minute epoxy is allowed to dry for at least 20 minutes, then any excess glue or smudges are removed using Kim wipes and a small amount of acetone and/or ethanol. (i) Two 1” lengths of flexible tubing are used as connectors to link the tubing of the exit ports to the tubing leading to the syringes. The 200 μL tips in the entry ports are dislodged from the glue with a firm twist and removed. (j) New 200 μL tips containing solution can be inserted into the entry ports and the solution can be drawn into the sample chamber by pulling suction on the syringe connected to the exit port.

In many cases, it is desired to load solutions into the sample chamber without removing the prism. Traditionally, this issue has been solved by gluing 200 μL tips and tubing directly into the holes of the slide [6]. However, we’ve found the tips can often come loose with this method, and if multiple samples are to be run through the same chamber, the repeated use of the entry tubing can cause contamination between samples. With this in mind, we have designed 3D printable ports which can be printed using a relatively cheap 3D printer (~$200 USD). The ports can be glued to the slide and support the insertion of a pipette tip directly into the holes of the sample chamber (figure 6 J-Q) (supplemental figure 11). The port on one end of the chamber is used to permanently secure a make-shift adapter into the hole of the slide (figure 6K), which is connected to syringes via FPLC tubing. The opposite port is used to support a 200 μL pipette tip which serves as a replaceable reservoir for solutions that can be drawn into the chamber via suction from the syringes (figure 6J, supplemental figure 11C). The benefit of this is that the incoming sample does not come into contact with tubing or other surfaces that might contain contaminates. Additionally, replacing the reservoir tip is as easy as drawing the solution into a fresh tip with a pipettor, inserting the new tip into the port, and ejecting the tip from the pipettor (supplemental figure 11B). Alternatively, the reservoir can be topped off by pipetting directly into the top of the existing reservoir tip and pulling the solution into the chamber using the syringe (supplemental figure 11A). Overall, these ports allow for multiple experiments to be conducted on a slide without the need to disassemble the prism, allow for multiple solutions to be flowed through the chamber during recording of a video without concern of sample contamination, and prevent the need to clean or replace tubing and syringes between experiments. The files necessary to print these ports are provided in the supplemental files.

Recording data from an experimental sample chamber is similar to recording a mapping file. However, extra care should be taken to minimize light exposure to the sample until ready to record. With that in mind, if the 3D ports are being used, it’s recommended to acquire TIR on a chamber filled with buffer before adding the fluorescent sample. After TIR is acquired, the sample can be loaded into the reservoir tip, and the recording can be started before or after the sample is pulled into the sample chamber with the syringe [21]. When the experimental video has finished recording, the video is processed using an environment such as IDL, which can operate scripts to convert the experimental video into a collection of trajectories [21, 31]. Each trajectory displays the fluorescence intensity of each emission channel observed at an immobilized molecule over the course of the video. Finally, the individual binding events seen in the trajectories are quantified, and a statistical approach is taken to analyze the various populations of event types and durations [9]. This can be accomplished using analysis tools such as the kinetic event resolving algorithm (KERA) provided by the Spies group (https://github.com/MSpiesLab/KERA), which can organize and sort the fluorescent trajectories from a range of single-molecule experiments [9, 35].

## 5. Applications

One of the primary reasons that single-molecule TIRF microscopy is becoming increasingly popular is due to its wide variety of applications, including the ability to study macromolecular complexes. Here, we briefly review select examples of how TIRF microscopy has been applied to the study of complex DNA structures and multi-component protein-DNA complexes responsible for homologous recombination, DNA replication, DNA repair, and transcription. Due to space constraints, we were limited in the number of investigators we could highlight in this section and apologize to those who have contributed to the development and application of single-molecule assays but were not mentioned herein. Specific techniques that utilize TIRF microscopes to study protein-DNA complexes include DNA or protein tethering [36–39], DNA combing and curtains [40–43], DNA tightropes (via oblique angle fluorescence imaging) [44, 45], as well as smFRET [37, 46–50] and TIRF/AFM systems [51–53]. While the examples mentioned below include data collected on both prism and objective based TIRF instruments, in most cases the prismTIRF microscope described above can be used for similarly designed experiments.

Much of the development and application of single molecule techniques suitable for the study of nucleic acid systems was pioneered by the lab of Taekjip Ha [54–64]. This includes the first demonstration of a protein diffusing on single-stranded DNA (ssDNA) [56], combining force and fluorescence to probe Holliday junction dynamics [55], exploring the activities of various DNA repair helicases [36, 58, 59, 61, 62], DNA strand exchange [63, 64], and nucleosome dynamics [60] to name a few. Importantly, this work laid the foundation for many other groups to further the development of single molecule techniques and explore other nucleic acid systems including recombination, DNA replication, DNA repair, and transcription.

Homologous recombination (HR) is an important mechanism for the repair of double-strand breaks (DSBs) and has been studied extensively via single-molecule techniques [40, 65–73]. This includes the ssDNA curtain assay developed by the Greene lab, which has provided new insight into HR regulation and the complex and heterogenous reaction intermediates that are involved [40, 73]. One example is the multi-functional Mre11/Rad50/Nbs1 (MRN) complex, which initiates DSB repair by recognizing and resecting the free DNA ends at damaged sites. The Finkelstein lab used DNA curtains to investigate how the MRN complex initiates DNA resection, assembles the larger resectosome, and is regulated by DNA-dependent protein kinase [65–67]. Another aspect of HR that has been studied using a TIRF microscope is the assembly of Rad51 filaments along ssDNA. Utilizing Protein Induced Fluorescence Enhancement (PIFE), which utilizes a fluorophore attached to the substrate as a reporter of the protein binding and dynamics [69], the Sua Myong lab monitored Rad51 filament formation on fluorescently labeled ssDNA substrates [70]. Interestingly, these studies suggest that filament formation occurs in the 5’ to 3’ direction and that the anti-recombinase Srs2 acts to transiently counterbalance Rad51 filament formation. Moreover, the Spies lab has used TIRF microscopes to follow the formation kinetics of filament nuclei at a single-monomer resolution, observing that human Rad51 nucleates and grows in dimers [71], which is in contrast to the addition/dissociation of monomers of *E. coli* RecA [72] and yeast Rad51 [70].

DNA replication requires several multi-factor components (i.e. a helicase, multiple DNA polymerases, a primase, and a ring-shaped sliding clamp protein). Because of the rapid and uneven fashion by which replisomes travel on DNA, several key aspects of their behavior are undetectable by ensemble studies. This has led multiple labs to use single-molecule fluorescence techniques to study the replisome [74–82]. For example, the van Oijen group has revealed new insight into the coordination of the bacteriophage T7 replisome by simultaneously monitoring the kinetics of DNA loop growth in the lagging strand and leading strand synthesis [76]. Around the same time, the Kowalczykowski lab used single-molecule analysis to determine that the leading and lagging strand polymerases function autonomously within a replisome and that replication is kinetically discontinuous and punctuated by pauses and rate-switches [77]. Additionally, the O’Donnell group was able to measure the processivity and rate of replisome activity in real-time using a single-molecule TIRF rolling circle replication assay, revealing that the lagging strand increases replisome processivity but slows replication fork progression [74]. Single-molecule TIRF has also been used to investigate how the structurally isolated *pol* and *5’-nuc* DNA binding domains of DNA polymerase I (Pol I) are utilized during interactions with DNA. Using smFRET, the Millar group identified separate subpopulations of DNA engaging the two DNA binding domains and found the relative populations and dwell times of each complex varied according to the nature of the DNA substrate [75]. Interestingly, additional experiments showed that DNA substrates can switch between the *pol* and *5’ nuc* DNA binding domains during a single encounter with Pol I.

DNA repair is usually accomplished by large multi-component repair complexes. One such component is Replication Protein A (RPA), a ssDNA binding protein which is required for many DNA repair pathways including nucleotide excision repair (NER), recombinational repair (HR and MMHJ), and mismatch repair (MMR) [38]. The Wold group applied single-molecule TIRF microscopy to analyze binding between DNA and surface-tethered RPA, revealing that RPA-DNA interactions can form two distinct complexes which are both critical for RPA function [38]. Single-molecule studies have also shown that RPA can resolve noncanonical G-quadruplex DNA structures which can interfere with DNA repair [83, 84]. To study another DNA repair factor, XPD helicase, the Spies group reported a single-molecule imaging strategy that utilized the FeS-mediated quenching of a site-specific fluorophore, which allowed for real-time and direct correlation between nanometer-scale domain motions and its interaction with individual DNA substrates [10]. Also using single-molecule TIRF, the Washington laboratory examined the assembly and disassembly of the multi-protein complex that carries out translesion synthesis (TLS) [85]. They found that ternary complexes containing proliferating cell nuclear antigen (PCNA) and two specialized TLS DNA polymerases, Rev1 and DNA polymerase η, have two architectures: PCNA tool belts and Rev1 bridges. Moreover, these complexes were shown to be dynamic and their architectures can interconvert without dissociation, possibly facilitating selection of the appropriate polymerase and polymerase-switching events during TLS.

Chromatin fibers are a key determinant of genome regulation as it dictates the accessibility of DNA to proteins. Single-molecule techniques have been instrumental in understanding protein-chromatin interactions [86–90]. For example, single molecule analysis has demonstrated that the nucleosome remodeler SWR1 destabilizes the DNA wrapped around the histone core, and this partial unwrapping is regulated by ATP binding [89]. The Fierz group used smFRET to directionally map local chromatin structural states and measure their interconversion dynamics [87]. In addition, by moving the fluorescent labels to several positions, structural information was obtained from several vantage points. This revealed that nucleosomes engage in effector protein regulated stacking interactions, which rapidly interchange on a timescale of micro-to-milliseconds. An additional study by the Lenstra lab combined *in vitro* and *in vivo* single-molecule imaging approaches, allowing them to observe the direct correlation between binding of the Gal4 transcription factor and the transcriptional bursting kinetics of the Gal4 target genes GAL3 and GAL10 in living yeast cells [86]. Interestingly, they found that the Gal4 dwell time sets the transcriptional burst size, and that the Gal4 dwell time is reduced by nucleosomes.

## Acknowledgements

We acknowledge the many groups that have made important contributions to single molecule studies which we were unable to highlight due to space limitations. This article was supported by National Institutes of Health (R01-ES029203 and R35-GM128562 to B.D.F) and (R35-GM131704 to M.S. and F.E.B), (K99-ES031148 to A.M.W.), and the Madison and Lila Self Graduate Fellowship to M.S.F.. We would also like to thank the Washington lab at the University of Iowa for providing training and technical support in the early stages of this build.

## Notes

### Competing Interest Statement

The authors have declared no competing interest.

