## Supplemental Figures for "Construction of a 3-color prism-based TIRF microscope to study the interactions and dynamics of macromolecules"

| Ref # | Item | Notes | Catalog ID | Vendor | Quantity |
|---|---|---|---|---|---|
| | Table | | | | |
| #1 | Optical Table | 3' x 6' x 12", 1/4-20 threads | RPR-36-12 | Newport | 1 |
| #2 | Table Legs | SL-600 Series, 22 in. height, 2400 lb max. capacity | SL-600-422 | Newport | 4 |
| #3 | Overhead Table Shelf | Fits 6 ft. Table, w/ electrical outlets | ATS-6 | Newport | 1 |
| | Mounting Hardware, Clamps, and Posts | | | | |
| #4 | 2" Post | 0.5" diameter, top tapped 8-32, bottom tapped 1/4-20 | 9621 | Newport | 30 |
| #5 | 3" Post | 0.5" diameter, top tapped 8-32, bottom tapped 1/4-20 | 9622 | Newport | 30 |
| #6 | 4" Post | 0.5" diameter, top tapped 8-32, bottom tapped 1/4-20 | 9623 | Newport | 30 |
| #7 | 6" Post | 0.5" diameter, top tapped 8-32, bottom tapped 1/4-20 | 9624 | Newport | 35 |
| #8 | 4" Adjustable Post Holder | Fits 0.5" diameter posts, bottom tapped 1/4-20 | VPH-4PK | Newport | 30 |
| #9 | Pedestal Base Adaptor | 1.25" diameter, 0.19" height, top thread 1/4-20 | PS-A-PK | Newport | 30 |
| #10 | Slotted Clamping Fork | Fits Pedestal Base Adaptor (#19), slot for 1/4-20 bolt | PS-F-PK | Newport | 30 |
| #11 | L-shaped Table Clamp | (for mounting #46, #20, #30) | CL5-P5 | Thorlabs | 20 |
| #12 | Right-Angle Post Clamp | Fits 0.5" diameter posts (#4-7) | 9935 | Newport | 15 |
| #13 | 1" Ultima Clear Edge Mirror Mount | Fits 1" diameter mirrors (#27-#29), 2 locking knobs, left-handed | U100-A-LH-2K | Newport | 15 |
| #14 | 0.5" Lens Mount | Fits 0.5" diameter lenses (#24-26, #50), 8-32 Thread | LH-0.5A | Newport | 4 |
| #15 | 1" Adjustable Post Holder + Pedestal Base | Fits 0.5" diameter posts (for stage mirror assembly) | VPH-1-P | Newport | 1 |
| | Excitation Lasers | | | | |
| #16 | OBIS 488nm LS 100mW Laser | CW, Diode, beam diameter 0.7±0.05 mm, (to excite AF488) | 1226419 | Coherent | 1 |
| #17 | OBIS 532nm LS 100mW Laser | CW, Diode, beam diameter 0.7±0.05 mm, (to excite Cy3) | 1261781 | Coherent | 1 |
| #18 | OBIS 640nm LX 100mW Laser | CW, Diode, beam diameter 0.8±0.1 mm, (to excite Cy5) | 1185055 | Coherent | 1 |
| #19 | OBIS Laser Heat Sink | Fits LX/LS OBIS Lasers, 2.7" height (optional) | 1193289 | Coherent | 3 |
| #20 | Vertical Translation Stage | 84 mm - 184 mm height, M6 thread holes (mounted via #11) | 860-0075 | Eksma | 3 |
| #21 | OBIS Laser Remote and Power Supply | Includes six 1-meter SDR cables | 1234466 | Coherent | 1 |
| | Laser Shutters | | | | |
| #22 | Laser Shutter | 3mm laser shutter, teflon coating, no electronic sync | LS3S2T0-100 | Uniblitz | 3 |
| #23 | Shutter Driver | four-channels, includes 710C shutter interconnect cables | VMM-D4 | Uniblitz | 1 |
| | Excitation Dichroic Mirrors and Filters | | | | |
| #24 | 488 nm Laser clean-up filter | 0.5" diameter, 488 nm, MaxLine® | LL01-488-12.5 | Semrock | 1 |
| #25 | 532 nm Laser clean-up filter | 0.5" diameter, 532 nm, MaxLine® | LL01-532-12.5 | Semrock | 1 |
| #26 | 640 nm Laser clean-up filter | 0.5" diameter, 640/8 nm, MaxDiode™ | LD01-640/8-12.5 | Semrock | 1 |
| #27 | Broadband Mirror | 1" diameter, 6.0 mm thick, RWE: λ/10 at 633 nm, 400 - 750 nm | BB1-E02 | Thorlabs | 15 |
| #28 | 505 nm Cutoff Longpass Dichroic Mirror | 1" diameter, 3.2 mm thick, TWE: λ/4 @ 633 nm | DMLP505 | Thorlabs | 1 |
| #29 | 550 nm Cutoff Longpass Dichroic Mirror | 1" diameter, 3.2 mm thick, TWE: λ/4 @ 633 nm | DMLP550 | Thorlabs | 1 |
| | Inverted Microscope Components | | | | |
| #30 | IX73 Microscope Frame (1 deck) | Single deck | IX73P1F | Olympus | 1 |
| #31 | 60X Water Objective (NA 1.20) | UPLSAPO60XW; U Plan S-Apo, WD0.28,W/CC0.13-0.21 | 1-U2B893 | Olympus | 1 |
| #32 | C-Mount Camera Adapter | Centerable (U-TV1XC) (1x Mag) | U-V111C | Olympus | 1 |
| #33 | Fluorescent Turret | (IX3-RFACS-1-2 CODED) | 5-UR416-1 | Olympus | 1 |
| #34 | Binocular Observation Tube | GX/IX (U-BI90-1-2) | 3-U243 | Olympus | 1 |
| #35 | 10X Eyepiece | (FN:22 WHN10X-1-7) | 2-U1007 | Olympus | 1 |
| #36 | 10X Eyepiece (Adjustable Focus) | (FN:22 WHN10X-H-1-7) | 2-U100H6 | Olympus | 1 |
| #37 | Left Handle Stage with Short Stalk | (IX3-SVL) | 4-U222 | Olympus | 1 |
| #38 | Stage Clips | FOR IX STAGE (IX-SCL) (includes 2) | FV4-U291 | Olympus | 1 |
| #39 | 6-position Nosepiece | (IX3-D6RES CODED) | U-R380 | Olympus | 1 |
| | Custom Stage Mounts and Prism Adapters | | | | |
| #40 | Custom Stage Adapter (Rear) | Schematics can be found in supplemental materials | CAD file ID 40 | machine shop | 1 |
| #41 | Custom Stage Adapter (Front) | Schematics can be found in supplemental materials | CAD file ID 41 | machine shop | 1 |
| #42 | Custom Stage Breadboard | Schematics can be found in supplemental materials | CAD file ID 42 | machine shop | 1 |
| #43 | Custom Prism Adapter | Schematics can be found in supplemental materials | CAD file ID 43 | machine shop | 1 |
| #44 | Custom Prism Connector Piece | Schematics can be found in supplemental materials | CAD file ID 44 | machine shop | 1 |
| #45 | Custom Prism Overhead Arm | Schematics can be found in supplemental materials | CAD file ID 45 | machine shop | 1 |
| #46 | Custom Camera Mount | Schematics can be found in supplemental materials | CAD file ID 46 | machine shop | 1 |
| #47 | Custom Slide Holder Insert | Schematics can be found in supplemental materials | CAD file ID 47 | machine shop | 1 |
| #48 | Finger Screws | ¼-20 thread, 5/8" long | 5PY63 | Grainger | 3 |
| #49 | Prism Adapter Bolts | 3/32" hex, 43494-N | 43494N | True Value | 3 |
| | Focusing Lens Micrometer Components | | | | |
| #50 | Plano-Convex Lens | 0.5" diameter, 50.0 mm focal length, uncoated | LA1213 - N-BK7 | Thorlabs | 1 |
| #51 | Optical Breadboard (4" x 6") | 1/4-20 thread on 1" grid, aluminum | SA2-04x06 | Newport | 1 |
| #52 | XYZ Quick-Mount Linear Stage | 0.5" travel, right-handed, ¼-20 thread | 460A-XYZ | Newport | 1 |
| #53 | Micrometer (13 mm Travel) | 1 µm vernier, 13 mm travel, 50.8 TPI (fits #52) | SM-13 | Newport | 3 |
| | Emission Dichroic Mirrors and Filters | | | | |
| #54 | AF488 Bandpass Filter | 1" diameter, 512/25, BrightLine® | FF01-512/25-25 | Semrock | 1 |
| #55 | Cy3 Bandpass Filter | 1" diameter, 585/40, BrightLine® | FF01-585/40-25 | Semrock | 1 |
| #56 | Cy5 Bandpass Filter | 1" diameter, 680/42, BrightLine® | FF01-680/42-25 | Semrock | 1 |
| #57 | 532 nm Cutoff Longpass Dichroic Mirror | 25.2 x 35.6 x 1.1mm, RWE: 1λ P-V @ 632.8 nm, BrightLine® | DI03-R532-T1-25X36 | Semrock | 1 |
| #58 | 635 nm Cutoff Longpass Dichroic Mirror | 25.2 x 35.6 x 1.1mm, RWE: 1λ P-V @ 632.8 nm, BrightLine® | DI03-R635-T1-25X36 | Semrock | 1 |
| | Camera and Emission Splitter | | | | |
| #59 | IXON ULTRA 897 EMCCD | 56 FPS, 512 X 512, 16 UM, USB | OAT-DU-897U-CS0-#BV | Olympus | 1 |
| #60 | Optosplit-III | Three-way image splitter, LS, 1X mag (contains #54-58) | O89-P280/310/0LS | Cairn-Research | 1 |
| | Tools and Screws | | | | |
| #61 | Hex Driver Set (Imperial) | 20-Piece Balldriver and Hex Key Kit, w/ Stand, Imperial | TC2 | Thorlabs | 1 |
| #62 | Hex Driver Set (Metric) | 15-Piece Balldriver and Hex Key Kit, w/ Stand, Metric | TC3/M | Thorlabs | 1 |
| #63 | 1/2" Spanner Wrench | 0.5" Spanner Wrench for Threaded Retaining Rings | LT05-WR | Newport | 1 |
| #64 | Imperial Screw and Hardware Kit (1/4"-20) | ¼-20 Cap Screw and Hardware Kit | HW-KIT2 | Thorlabs | 1 |
| #65 | Metric Screw Kit (Metric M4-M8) | Metric Screw Kit | 12980 | Precision | 1 |
| #66 | Front Stage Breadboard Bolts | #10-24, 1" long, Philips | 3248-P | True Value | 2 |
| #67 | Back Stage Breadboard Bolts | #10-24, ¾" long, Hex | 3247-D | True Value | 2 |
| #68 | 1" Iris | 25.0 mm max aperture, TR3 Post (used for alignment) | ID25 | Thorlabs | 4 |
| #69 | Thin Slip-On Post Collar | Fits 0.5" diameter posts (used to maintain height of iris during use) | R2T | Thorlabs | 5 |
| #70 | Beam Height Measurement Tool | 12" tall, (used to measure height of beam and iris) | BHM4 | Thorlabs | 1 |
| | Miscelaneous / Consumables | | | | |
| #71 | Pellin-Broca Quartz Prism | Fused Silica | 325-1206 | Eksma | 3 |
| #72 | 5 Minute Epoxy | Devcon, syringe | 12084 | Tap Plastics | 1 |
| #73 | Quartz Microscope slide | 1" x 3" x 1mm | 1X3X1MM | Finkenbeiner, Inc. | 10 |
| #74 | TetraSpeck Microspheres | 0.2 µm diameter, fluorescent blue/green/orange/dark red | T7280 | Thermo Fisher | 1 |
| #75 | Plain Glass Microscope Slides | Glass, 25 x 75mm, 90° Ground Edges, Plain | 1301 | Globe Scientific | 144 |
| #76 | Low Autofluorescence Immersion Oil | n = 1.518, Olympus Type F, 30 mL | MOIL-30 | Thorlabs | 1 |
| #77 | Solvent Dropper Bottle | Solvent dropper bottle, 2 oz. (60ml) (to add water to objective) | LAB-14 | Newport | 1 |
| #78 | ETFE Tubing (ID 1.0 mm, OD 1/16") | 3 m length, translucent | 18114238 | Cytiva | 1 |
| #79 | Fisherbrand™ Redi-Tip™ 200uL Pipet Tips | General purpose, yellow, 1000/PK | 02-707-500 | Fisher Scientific | 1 |
| #80 | HEPA Air Purifier | Holmes small room 3-speed, w/ optional ionizer | B0000DK35B | Holmes (Amazon) | 1 |
| #81 | 3M Double-Sided Adhesive Sheet | Clear, 5 MIL, double linered, 12"X12" | 7955MP | Hisco | 100 |

Table 1. Component List.

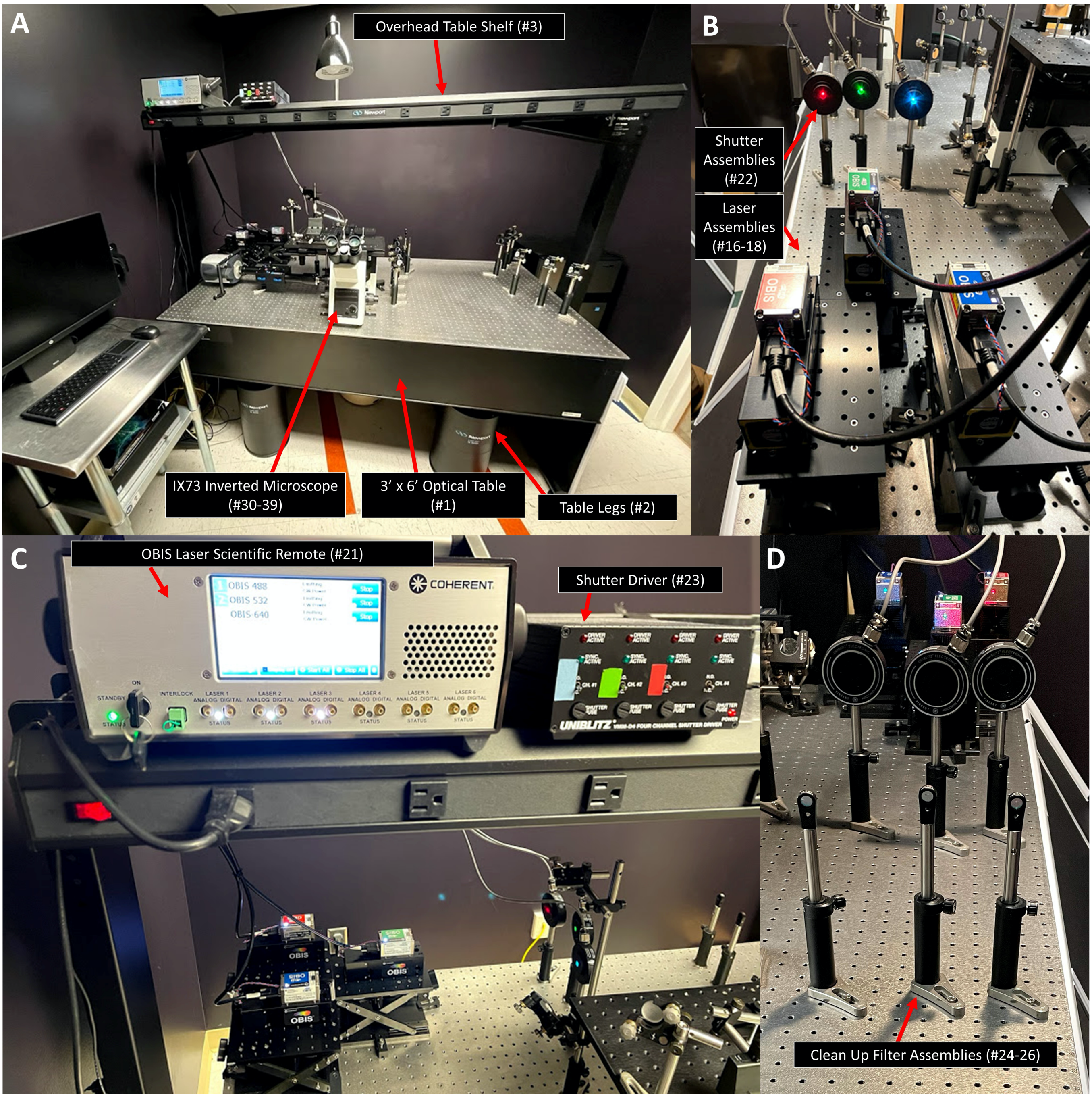


Supplemental Figure 1. Lasers, shutters, and clean up filters. (A) The prismTIRF microscope components are assembled on an optical table which is supported by four pneumatic legs. An overhead shelf is used to store the OBIS laser scientific remote and shutter driver. (B) Each laser head is mounted to a heat sink, secured to a vertical translation stage, and attached to the table using three L-shaped clamps. Each shutter is attached to a 6″ post, an adjustable post holder, a pedestal base adaptor, and secured to the table with a slotted clamping fork. (C) The OBIS laser scientific remote and shutter driver are positioned on the overhead shelf within reach of the operator. (D) Each laser clean-up filter is mounted inside a fixed lens mount and attached to a 6″ post, an adjustable post holder, a pedestal base adaptor, and secured to the table with a slotted clamping fork.

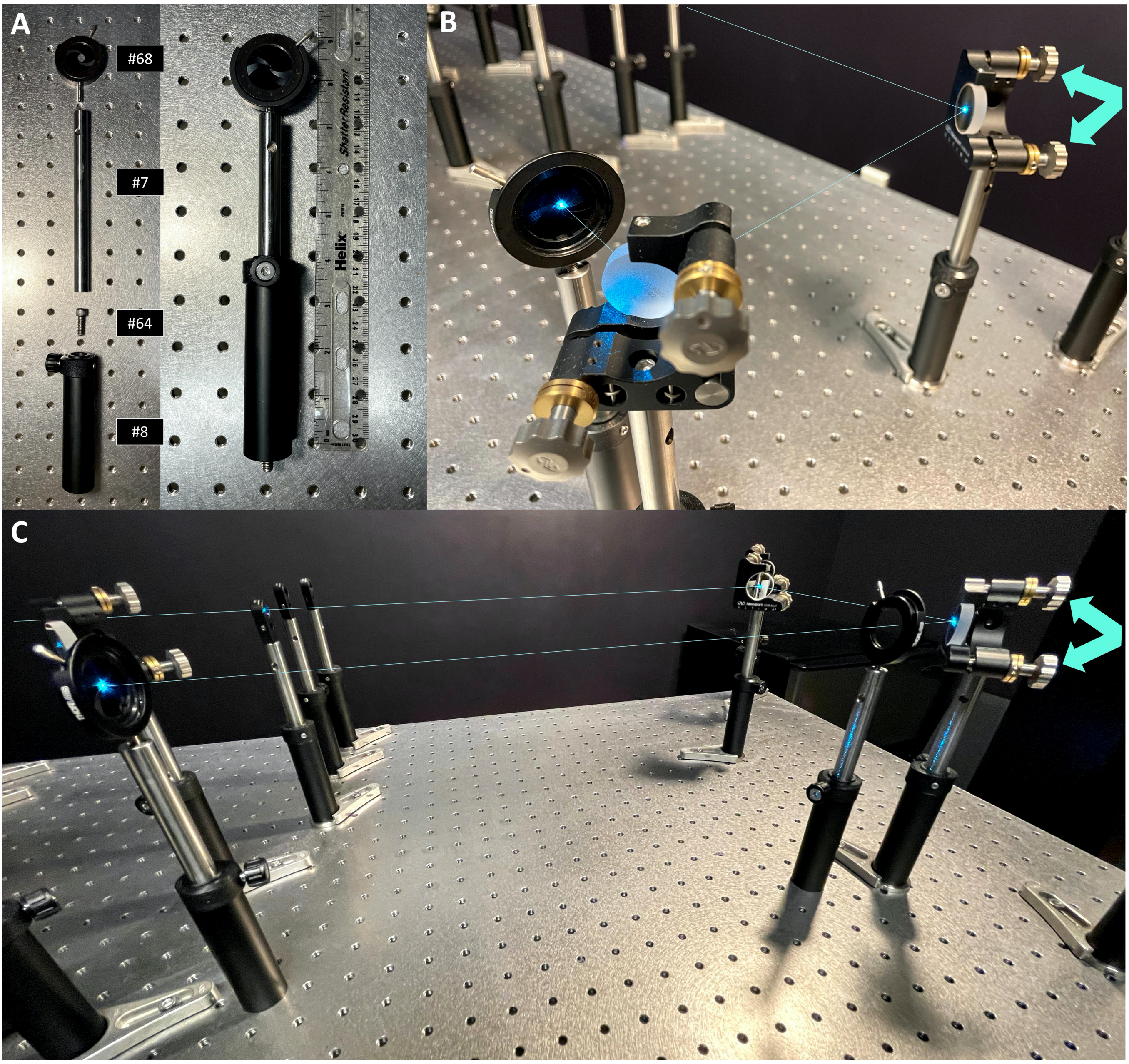


Supplemental Figure 2. Leveling the excitation beams. (A) Leveling beams using a two-mirror turn requires two irises. Each iris is attached to a 6″ post and an adjustable post holder. Before inserting the post into the post holder, a ¼-20 bolt is screwed into the bottom of the post holder so the post holder can be mounted directly into the holes of the optical table. To make an 8″ iris assembly, the post is adjusted within the post holder such that the center of the aperture is 8″ off the surface of the table. A slip-on post collar (table 1, #69) can be attached to allow for the post to be rotated within the post holder while maintaining the desired height (not shown). (B) To level a beam using a two-mirror turn, the aperture of the first iris is constricted and the adjustment knobs on the mount of the first mirror are tuned until the beam passes through the very center of the first iris. (C) Then, the aperture of the first iris is opened and the adjustment knobs on the mount of the second mirror are tuned until the beam passes through the very center of the second iris. A and B are repeated iteratively until the beam passes through the very center of both constricted irises.

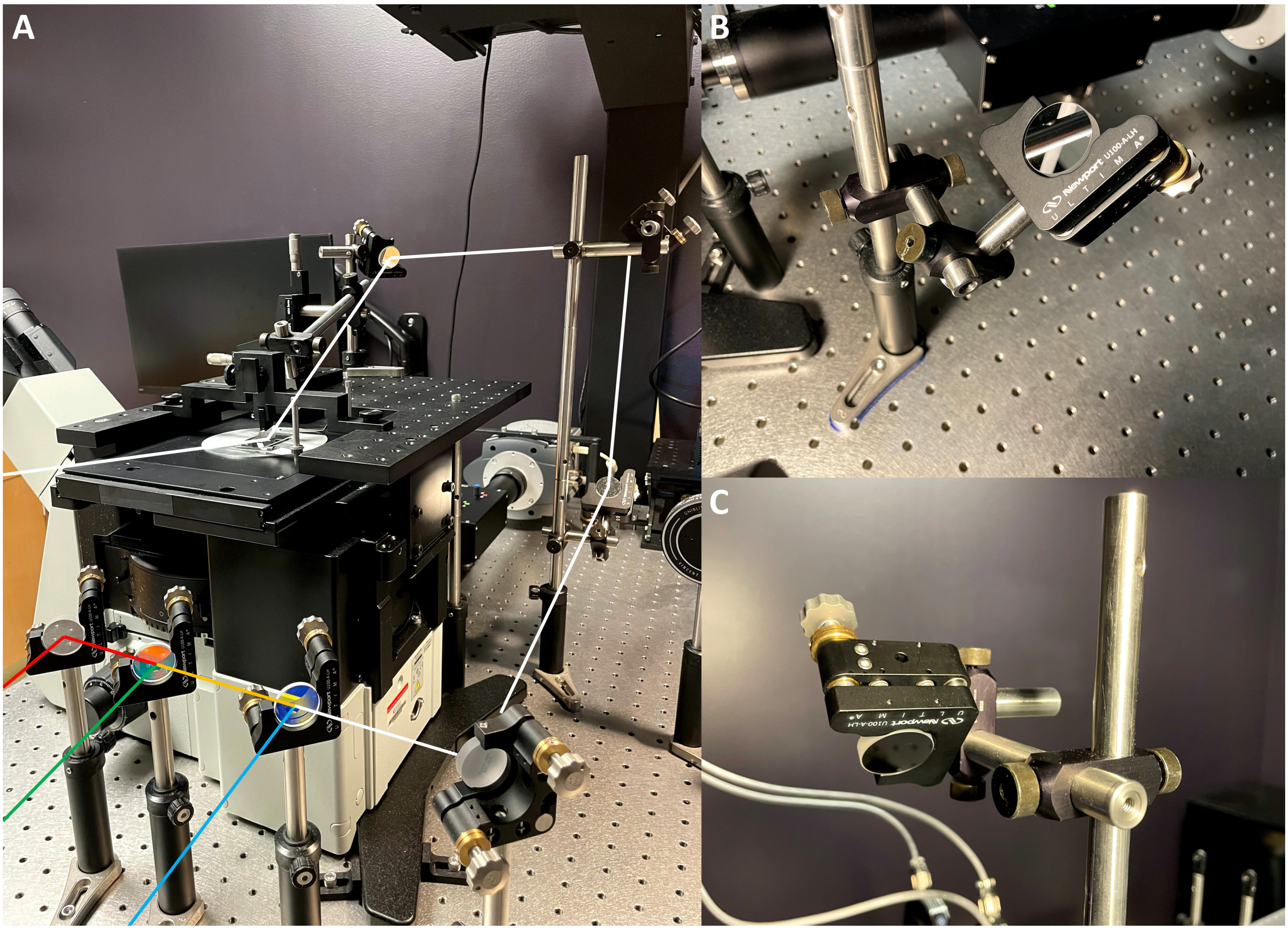


Supplemental Figure 3. Beam path to microscope stage. (A) After the beams are leveled and co-aligned, they are directed to the microscope stage using a series of broadband mirrors. (B-C) Two broadband mirrors attached to a periscope are used to raise the beam to the level of the microscope stage. The periscope is assembled from three 6″ optical posts, an adjustable post holder, a pedestal base adaptor, and secured to the table near the O26 hole with a slotted clamping fork. (B) The bottom broadband mirror is mounted inside an Ultima clear edge mirror mount, connected to a 2″ post, and attached to the 18″ post with two right angle clamps and a 3″ post. (C) The top broadband mirror is mounted inside an Ultima clear edge mirror mount, connected to a 2″ post, and attached to the 18″ post with two right angle clamps and a 4″ post. The top and bottom mirrors are adjusted iteratively until the beam is directed vertically by the bottom mirror and directed horizontally towards the stage by the top mirror.

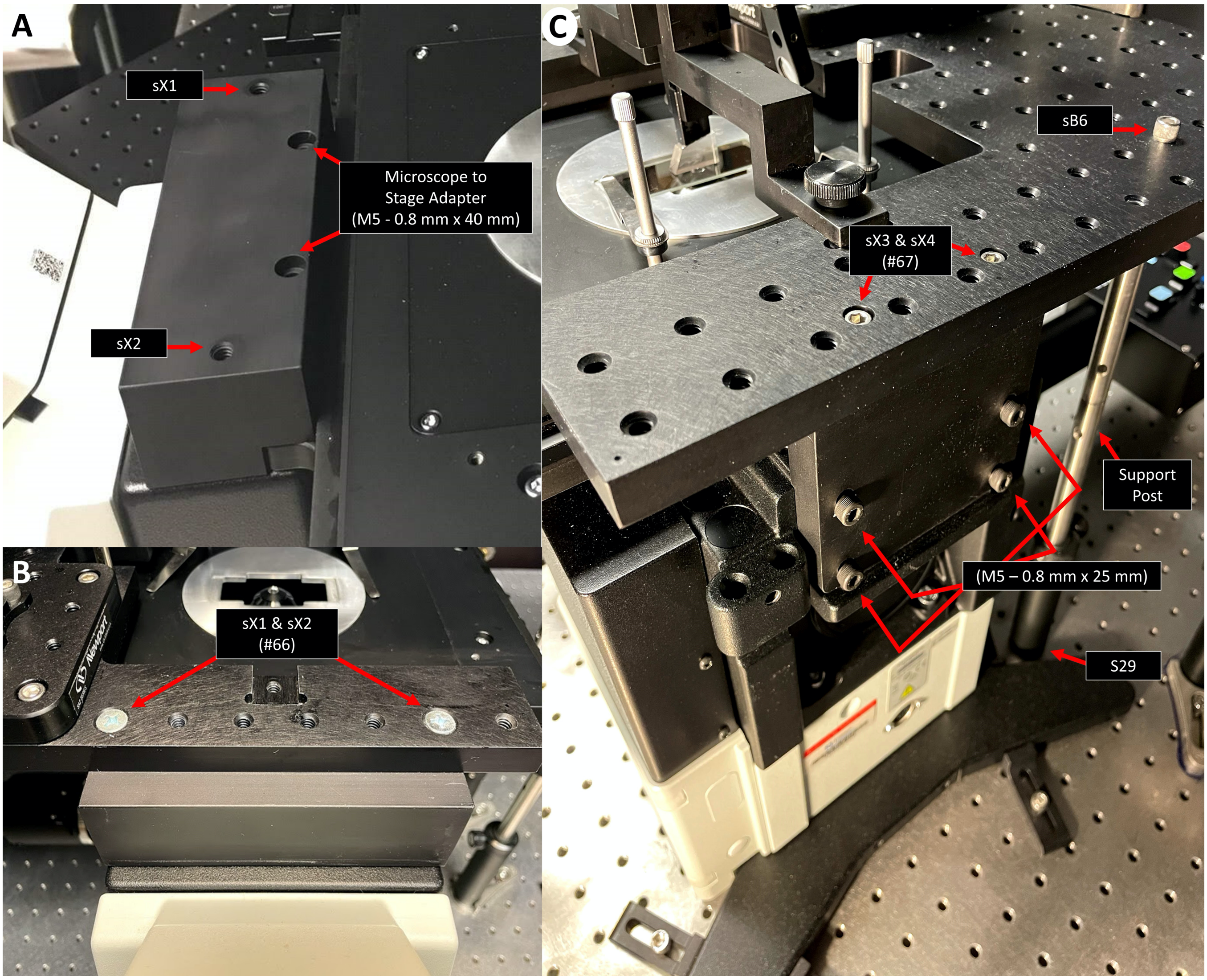


Supplemental Figure 4. Mounting the stage breadboard. (A) The stage breadboard is secured to the inverted microscope via two adaptor pieces. The front stage adapter is bolted to the microscope just behind the binocular tube using two 40 mm long M5 bolts. The stage breadboard is bolted to the front stage adapter through the sX1 and sX2 holes. (B) The rear stage adapter is bolted to the back of the inverted microscope using four 25 mm M5 bolts. The stage breadboard is then bolted to the rear stage adapter through the sX3 and sX4 holes using two 1″ long #10-24 bolts. A support post can be assembled from two 6″ posts, an adjustable post holder, and a pedestal base adapter and bolted to the stage breadboard through the sB6 hole using a 1/4″-20 bolt.

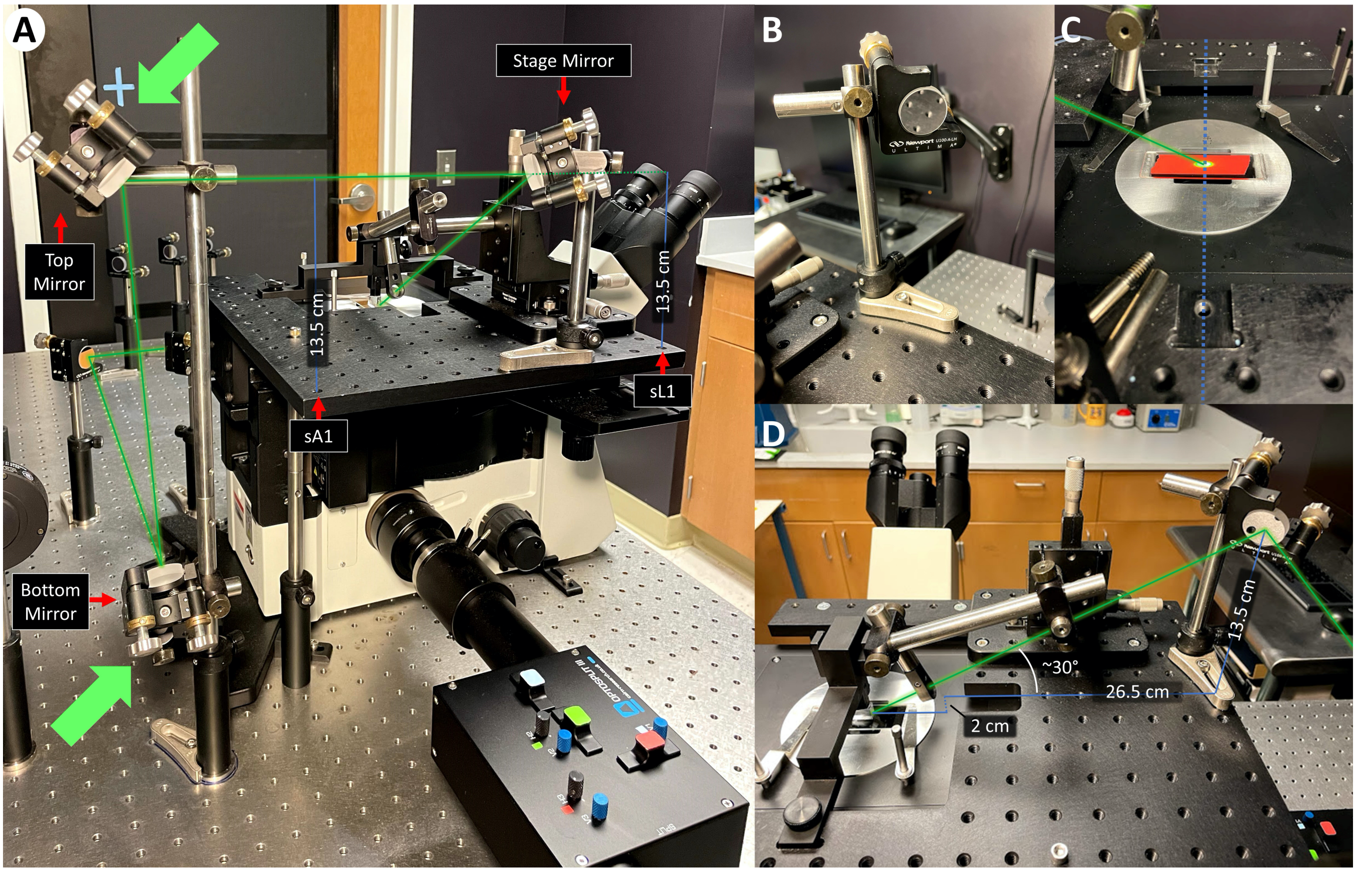


Supplemental Figure 5. Stage mirror and TIRF angle. (A) The top mirror of the periscope is used to direct the beam over the sA1-sF1 holes of the stage breadboard and onto the stage mirror. Before mounting the stage mirror to the stage breadboard, two 13.5 cm tall irises are mounted to the sA1 and sL1 holes of the stage breadboard and the top and bottom mirror are adjusted (green arrows) until the beam passes through the very center of both apertures. When finished, the irises are removed. (B) To assemble the stage mirror, a broadband mirror is inserted into a clear-edge mirror mount, attached a 3″ post, combined to a 6″ post using a right-angle clamp, and then secured to the stage breadboard using a 2″ post holder and a slotted clamping fork. (C) The stage mirror is positioned such that the beam strikes the face of the mirror directly above the sG1 hole, traverses over sG2-sG6, and strikes the microscope stage between the overhead bar mounting holes (sX5 and sX6). (D) The appropriate height at which the beam should strike the stage mirror is dependent on the horizontal distance between the prism and where the beam strikes the stage mirror. In the case of this set up, the horizontal distance between the prism and the reflection point on the stage mirror was measured to be approximately 26.5 cm. Additionally, the microscope stage is approximately 2 cm below the stage breadboard. Thus, the beam should strike the stage mirror 13.5 cm from the surface of the stage breadboard to induce the 30° incident angle at the prism face.

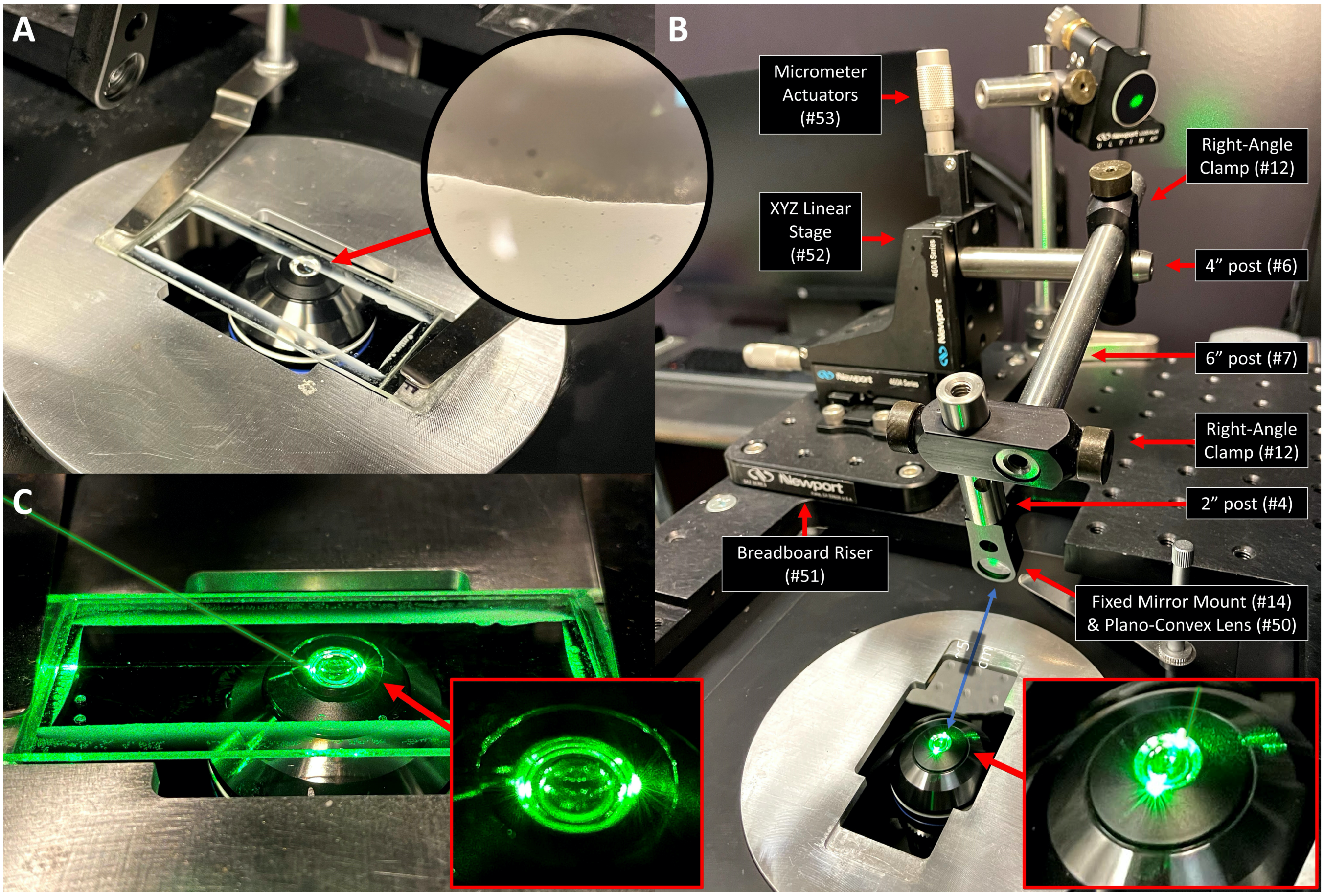


Supplemental Figure 6. Planco-convex lens and XYZ linear stage assembly. (A) The objective is used as a target while setting up the lens assembly and therefore needs to be positioned correctly before assembling the lens and XYZ linear stage. To position the objective, a bead slide is placed on the microscope stage and the inside edge of the chamber is brought into focus. When complete, the objective is at (or very close to) the correct position to observe TIRF within the chamber. (B) The lens assembly consists of a 4″ x 6″ breadboard riser, an XYZ linear stage, three micrometer actuators, two right-angle clamps, a 4″ post, a 6″ post, a 2″ post, a fixed mirror mount, and the plano-convex lens. After constructing the lens assembly and mounting it to the stage breadboard, the three micrometers are set to the center of their travel range (in this case ~6 mm). Then, the right-angle clamps and optical posts are adjusted until the lens is roughly 5 cm from the objective, with the beam passing through the center of the lens and striking the near edge of the objective's front lens. (C) When the lens is positioned properly, the beam will produce symmetric points of light on either side of the objective. 5mm on pic

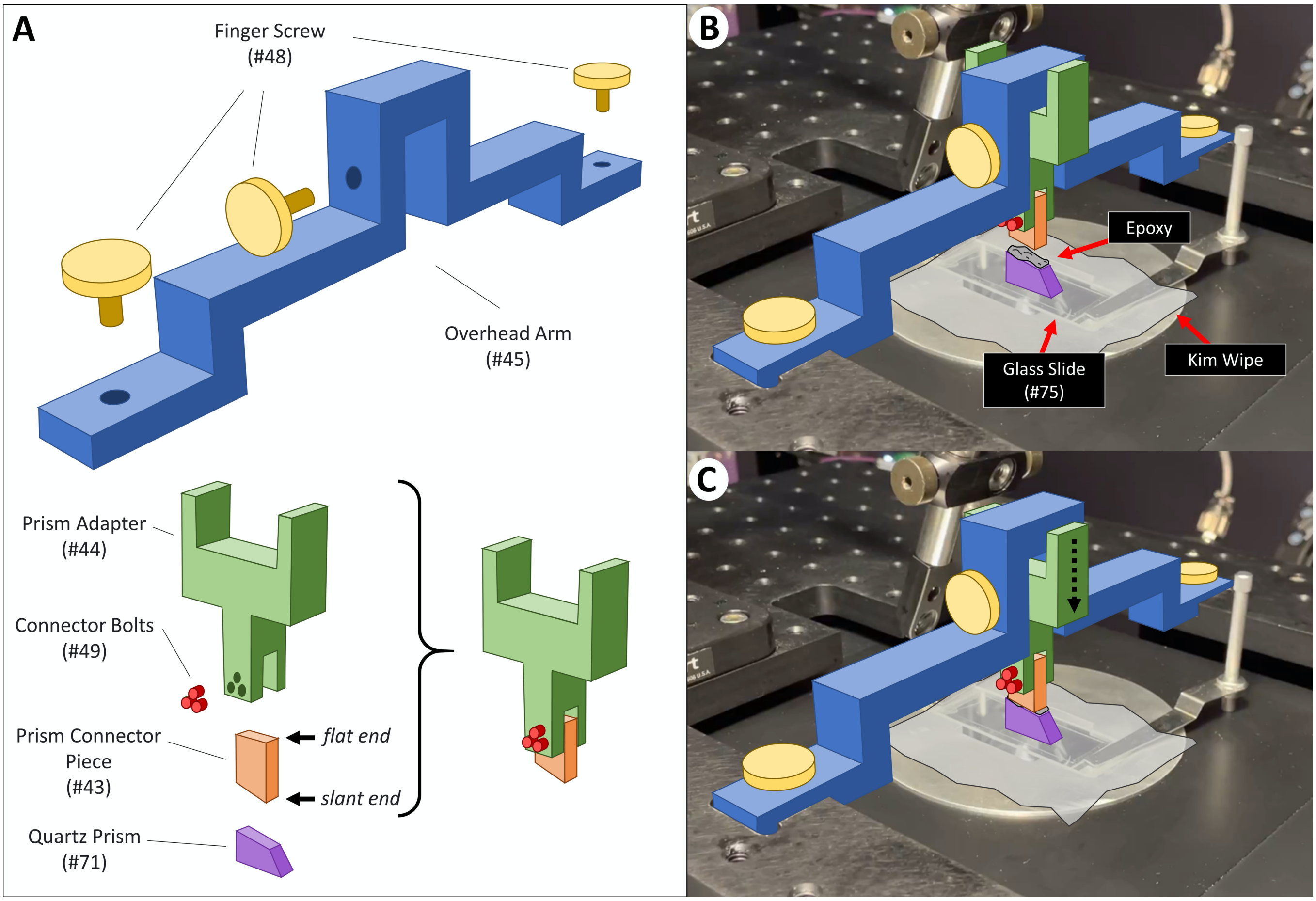


Supplemental Figure 7. Mounting the prism. (A) The prism assembly consists of a quartz prism (table 1, #71), a prism connector piece (table 1, #43), three connector bolts (table 1, #49), a prism adapter (table 1, #44), an overhead arm (table 1, #45), and three finger screws (table 1, #48). To build the prism assembly, the prism connector piece is first inserted into the prism adapter such that the flat end is facing the adapter. The slanted end of the prism connector piece is angled to match the contours of the prism. The prism connector piece is secured to the prism adapter by tightening the three connector bolts until they are snug, but not overtightened, which could cause the aluminum adapter to bend. (B) To attach the prism to the prism connector piece, a standard glass slide is first secured to the microscope stage using the stage clips and a Kimwipe is laid over the glass slide. Then, the overhead bar is attached to the stage breadboard using two finger screws at sX5 and sX6. The prism adapter is then inserted into the central nook of the overhead bar such that the three connector bolts face towards the sX6 hole, and the prism adapter is secured in place using the central finger screw. A smear of 5-minute epoxy is applied to the top of the prism, and the prism is placed on the Kimwipe directly under the prism connector piece. (C) The central finger screw is then loosened, and the prism adapter is carefully lowered until the prism connector piece makes contact with the top of the prism. The central finger screw is then tightened to secure the prism adapter into place. If needed, a pipette tip can be used to nudge the prism and straighten it with respect to the prism connector piece. The glue is allowed to set overnight. Over a period of use, the prism may become chipped or scratched. When this occurs, the prism can be snapped off of the prism connector and replaced with a new prism.

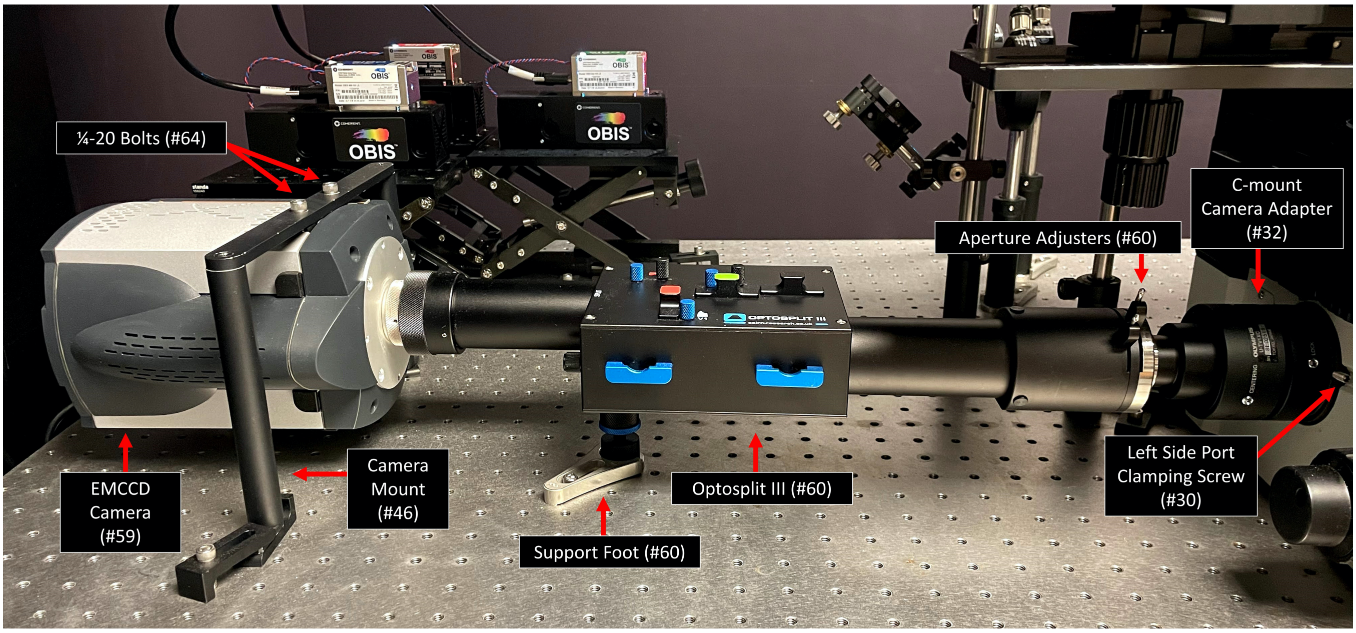


Supplemental Figure 8. Emission path. The emission light path is directed to the left side port of the inverted microscope where the emission from the three fluorophores will be split using the Optosplit III and projected onto the EMCCD Camera. To assemble the components within the emission path, the left side port cap is removed from the microscope by loosening the left side port clamping screw. Then, the C-mount camera adapter is secured to the entry port of the Optosplit III and inserted into left side port. The support foot of the Optosplit III is attached and adjusted such that the body of the Optosplit III is level with the plane of the table. The C-mount camera adapter is then secured in place by tightening the left side port clamping screw. The EMCCD camera is attached to the camera mount (table 1, #46) by removing the two rubber stoppers on the side of the EMCCD camera and using two ¼-20 bolts to secure the camera mount in place. The EMCCD camera can then be fixed onto the output port of the Optosplit III. If the EMCCD displays an image that is slanted, the bolts securing the camera to the camera mount can be loosened or tightened to adjust the camera orientation which will rotate the depicted image. In the configuration shown here, the camera is turned 90° with respect to the Optosplit III, which allows for the USB and power cables to protrude from the side instead of down towards the table where there is little space. Although this will result in the image appearing to be rotated 90° clockwise, it can be easily corrected by rotating the image 90° counterclockwise in the camera settings and/or collection software settings.

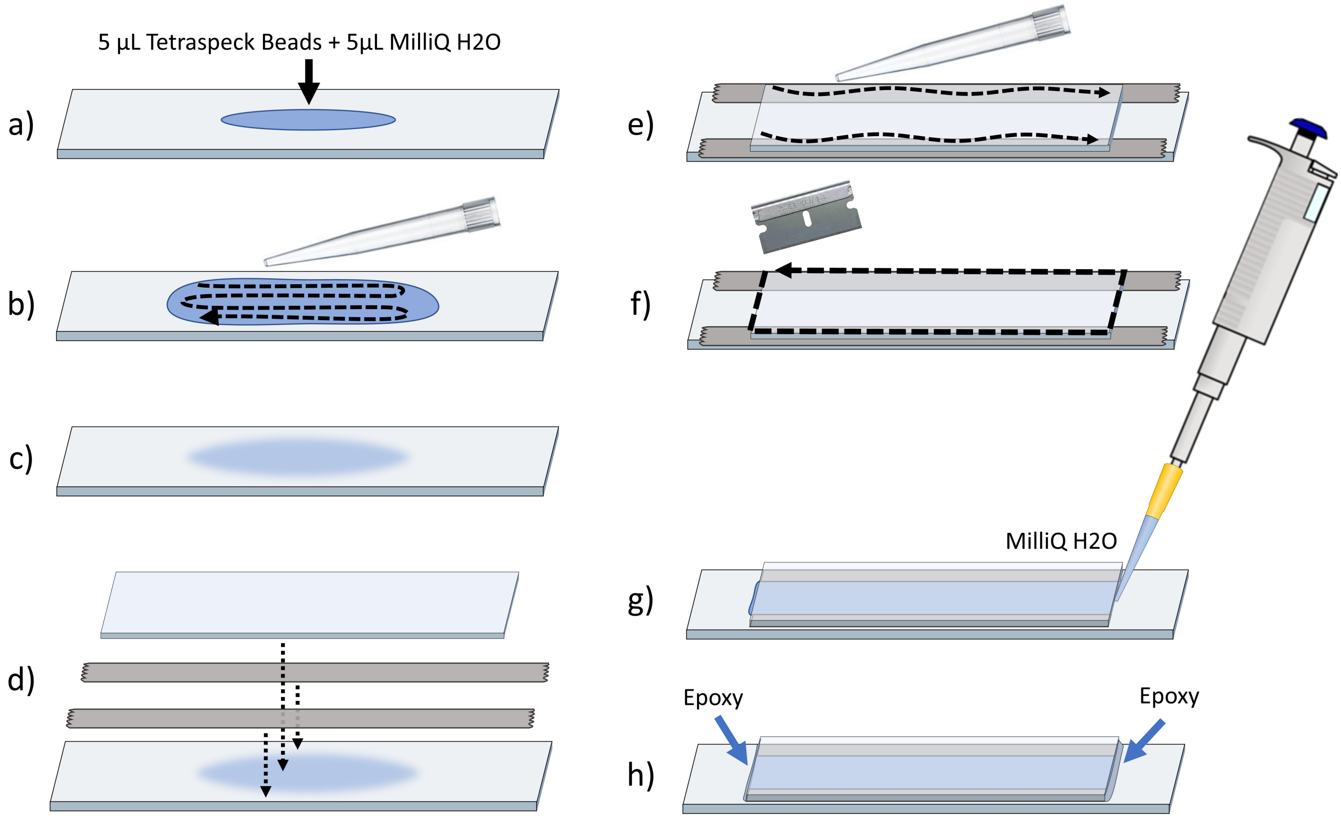


Supplemental Figure 9. Bead slide assembly. (A) 5 µL of concentrated (~2.3 × 10^10 particles/mL) 0.2 µm TetraSpeck™ fluorescent microspheres and 5 µL of MilliQ water are pipetted onto the center of a glass slide. (B) The 10 µL volume is spread over the central region of the slide using a pipette tip. (C) The slide is covered and allowed to dry. (D) Two pieces of double stick tape are placed across the long edges of the slide on either side of the dried beads and a coverslip is placed in the center of the glass slide, on top of the double stick tape. (E) A pipette tip is used to press on the coverslip and seal the bond between the coverslip and the tape. (F) The excess tape surrounding the coverslip is removed with a razor. (G) The space between the coverslip and the slide is filled with MilliQ water. (H) 5-minute epoxy is used to seal the short edges of the enclosure. After 20 minutes, the glue will have set and the slide can be used. A bead slide can last for over a month, but the image quality will eventually depreciate, and the slide will need to be replaced.

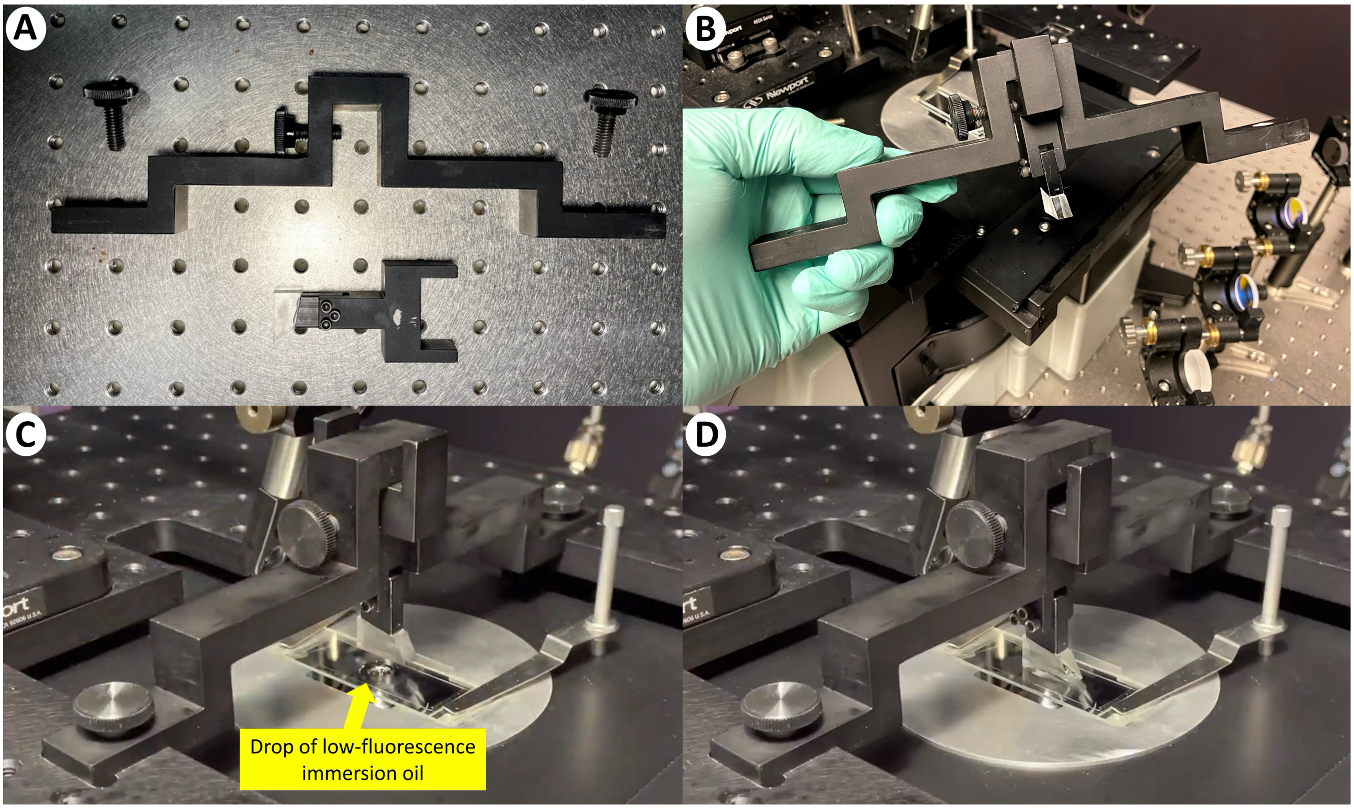


Supplemental Figure 10. Mounting the prism during operation. (A) The prism is held in place above the slide using the prism adapter and overhead arm. (B) To mount the prism assembly onto the microscope during operation, the prism adapter is first inserted all the way up into the central nook of the overhead bar and secured into place by tightening the central finger screw. (C) After securing the slide to the microscope stage with the stage clips and placing a drop of low autofluorescence immersion oil on the slide, the overhead bar is inserted into the slots of the stage breadboard and secured in place by tightening the finger screws at sX5 and sX6. (D) After the overhead bar is secured to the stage breadboard, the top of the prism adapter is held with one hand to prevent it from dropping, while the other hand loosens the central finger screw. With the prism adapter free, the prism is gently lowered onto the drop of oil and secured in place by tightening the central finger screw. To remove the prism assembly, these steps are performed in reverse.

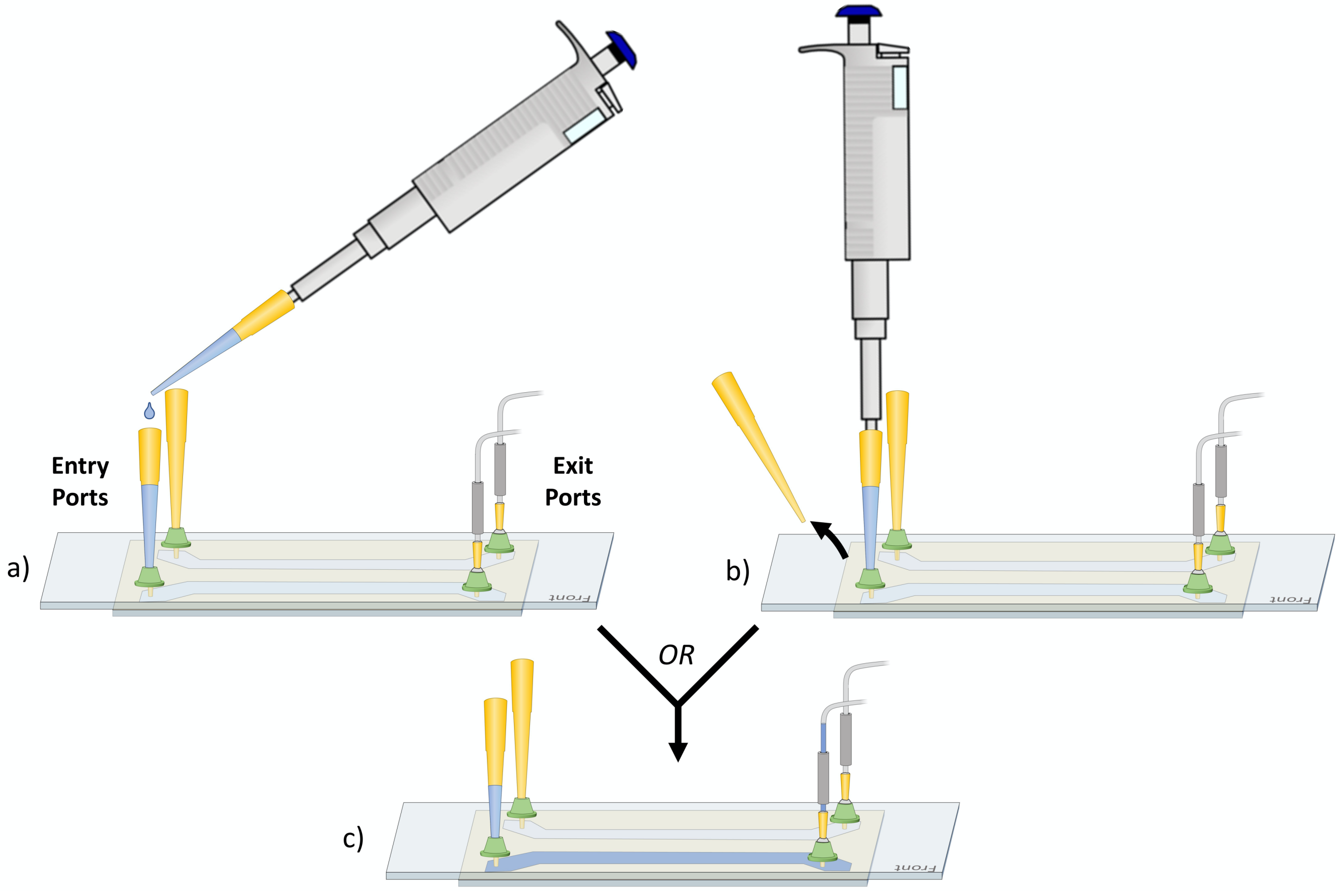


Supplemental Figure 11. Using a sample chamber. (A) To allow for easy and secure insertion of pipette tips into the holes of the sample chamber, we have designed 3D printable ports which can be glued to the surface of the slide. The entry ports located on one side of the chamber are used to support 200 μL pipette tips, which serve as replaceable reservoirs for solutions. The reservoirs can be filled by pipetting directly into the top of the pipette tip. (B) Alternatively, the reservoir can be replaced by removing the old tip from the entry port, pulling fresh solution into a new tip with a pipettor, inserting the tip into the empty entry port, and ejecting the tip from the pipettor. (C) The solution in the reservoir can then be pulled through the sample chamber by pulling negative pressure on the syringe attached to the exit port on the opposite side. Care should be taken to avoid flowing air through the sample chamber, which can cause issues with the immobilized substrates and/or fluorescent molecules within the chamber.
